# An axiomatic approach to cultivar ranking in multi-environment trials

**DOI:** 10.64898/2026.06.27.734959

**Authors:** Aleksei Y. Kondratev, Egor Ianovski, Ekaterina Voronina, José Crossa

**Affiliations:** HSE University, St. Petersburg, Russia; independent researcher, Auckland, New Zealand; CIMMYT Emeritus, International Maize and Wheat Improvement Center (CIMMYT), Texcoco, MexicoDe; Department of Statistics and Data Science, Post Graduate College, Montecillo, Texcoco, Edo. de Mexico, Mexico

**Keywords:** plant breeding, geometric mean, ranking rule, yield stability, Pareto efficiency, independence of irrelevant alternatives

## Abstract

Multi-environment trials are central to cultivar evaluation because they reveal how candidate cultivars perform across locations, years, management conditions, and stress environments. The resulting yield matrix is a rich source of data on genotype-by-environment interaction, and a wide literature on estimation, decomposition, visualisation, and prediction of yield potential and stability has flourished. However the ultimate question of which cultivar to recommend on the basis of such a matrix is often left implicit. The question is far from trivial, and in this paper we formulate cultivar recommendation as an axiomatic ranking problem. This framework is rich enough to encompass the existing literature on stability indices, as well as any other deterministic ranking procedure. We show that many commonly used stability-based procedures can violate minimal criteria of efficiency or consistency. The result of such violations is that a cultivar with uniformly high yield could be ranked below a cultivar with uniformly low yield, or the relative ranks of two cultivars could depend on whether or not a third cultivar is present in the matrix. Our results prove that under a small number of such criteria the space of admissible rules collapses to the family of power means and their limiting cases. If we further wish to allow multiplication normalisation of yield, we are left with the geometric mean as the unique solution.

## 1. Introduction

Plant breeding programs aim to identify and recommend cultivars that combine high productivity, acceptable stability, and reliable performance across the target population of environments. Their success has been indisputable. For well over a century, sustained genetic improvements on the order of 0.5–1% per year have been observed in maize, wheat, rice, soybean, and other crop systems, increasing both yield and resistance to adverse growing conditions (Morgounov et al., 2010; Di Matteo et al., 2016; Boehm Jr et al., 2019; Voss-Fels et al., 2019; Wang et al., 2021; Abdala et al., 2024; Gambin et al., 2025). As agriculture faces increasing pressure from population growth, climate change, land limitation, and input constraints, the identification and deployment of superior cultivars remains one of the most effective strategies for increasing food production without expanding cultivated land (Snowdon et al., 2021; Laidig et al., 2021; Senapati et al., 2022).

A central component of this process is the multi-environment trial. Candidate cultivars are evaluated across environments defined by locations, years, management practices, irrigation regimes, disease pressures, or stress conditions (Crossa, 1990; Yan and Kang, 2003; Smith et al., 2005; van Eeuwijk et al., 2016). The resulting data form a yield profile in which each cultivar is represented not by a single value, but by a vector of environment-specific yields. This structure reflects the biological and agronomic reality that cultivar yield is environment-dependent. A cultivar may perform exceptionally well in favourable environments but poorly under stress, whereas another cultivar may show moderate but consistent yield.

A large and sophisticated literature exists to analyse the outcomes of such trials. Early regression-based and parametric stability approaches, including the Finlay–Wilkinson regression, the Eberhart–Russell model, Shukla’s stability variance, and related statistics, interpreted stability through cultivar responses to environmental conditions, deviations from regression, or cultivar-specific contributions to genotype-by-environment interaction (Finlay and Wilkinson, 1963; Eberhart and Russell, 1966; Shukla, 1972). Later, AMMI models combined analysis of variance with principal component analysis to decompose genotype-by-environment interaction into interpretable multiplicative components (Gauch Jr, 1988; Zobel et al., 1988; Crossa, 1990). SREG and GGE biplot approaches provided graphical tools for visualising cultivar yield, identifying mega-environments, detecting crossover interaction, and comparing cultivars relative to environmental patterns (Yan et al., 2000; Yan and Kang, 2003). Related linear–bilinear and shifted multiplicative models have also been used to study crossover genotype-by-environment interaction, classify cultivars and environments, and identify patterns of specific adaptation not captured by simple mean-based comparisons (Cornelius et al., 1992; Crossa and Cornelius, 1997; Crossa et al., 2002, 2004).

Modern statistical methodology has further improved cultivar evaluation. Mixed models are now central because they can accommodate unbalanced trials, account for experimental design, model heterogeneous error variation, adjust for spatial field trends, and estimate genotype, environment, and genotype-by-environment effects within a unified framework (Piepho, 1997; Smith et al., 2001, 2005; Burgueño et al., 2011; Malosetti et al., 2016; van Eeuwijk et al., 2016). Factor-analytic models provide a parsimonious way to model heterogeneous genetic correlations among environments, thereby improving prediction and interpretation in large and unbalanced trial networks (Piepho, 1997; Smith et al., 2001, 2005, 2015). Genomic prediction models have extended cultivar evaluation by using genome-wide marker information, often together with pedigree, environmental covariates, or multi-environment covariance structures, to predict the yield of tested and untested cultivars across target environments (Meuwissen et al., 2001; Jannink et al., 2010; de los Campos et al., 2013; Jarquín et al., 2014; Crossa et al., 2017; Alemu et al., 2024). High-throughput phenotyping has further expanded the information available to breeding programs by enabling repeated, large-scale measurement of crop growth, stress response, and adaptation under field conditions (Cabrera-Bosquet et al., 2012; Araus and Cairns, 2014; Araus et al., 2018).

These approaches and technologies deal with the estimation, decomposition, visualisation, and prediction of genotype-by-environment interaction. They are essential for cultivar evaluation, but they do not by themselves define how cultivars should be finally ranked for recommendation. The breeder still faces a distinct decision problem: given two vectors of yield data for two different cultivars, how to determine which cultivar is better? Given *m* such vectors, how to rank the cultivars from most recommended to least?

This distinction is the starting point of the present paper. We do not propose another model for genotype-by-environment interaction or another descriptive stability statistic. Instead, we formulate cultivar recommendation in multi-environment trials as an axiomatic ranking problem. A ranking rule is a procedure that takes the yield profile and returns an ordered list of cultivars. The yield profile may contain not only observed yields, but also adjusted, estimated, or predicted yields from any appropriate statistical model. In this sense, the proposed framework is complementary to the vast literature on genotype-by-environment interaction. It begins after these methods have produced yield profiles and asks which ranking rules are logically defensible for final recommendation.

The need for such a framework arises because yield potential and yield stability are not always aligned. The naïve solution to rank cultivars by arithmetic mean yield favours high overall productivity but may insufficiently penalise cultivars with very poor performance in some environments. Conversely, there exists a large literature on metrics to measure the stability of cultivar yield (Mohammadi and Amri, 2008; Reckling et al., 2021; Pour-Aboughadareh et al., 2022), but these may favour cultivars with low variability but limited productivity. A cultivar may be stable simply because it performs poorly everywhere. A minimal requirement for a cultivar-ranking rule is that it should not prefer a uniformly inferior cultivar over a uniformly superior one (Mead et al., 1986). If one cultivar yields at least as much as another cultivar in every environment, then the superior cultivar should not be ranked below the inferior cultivar. This requirement is known as weak Pareto efficiency (Chebotarev and Shamis, 1998). It is simple, agronomically natural, and directly relevant to cultivar selection. Nevertheless, most commonly used stability-based procedures violate this requirement.

To illustrate this point consider Table 1, which contains the results of a hypothetical trial involving ten cultivars and five environments, and Table 2, which contains the stability rankings produced by sixteen common stability measures applied to Table 1. Observe that every measure ranks either *c*_3_, *c*_4_, *c*_5_, or *c*_6_ as the most stable, whereas the yield of *c*_1_ is better than all of *c*_3_, *c*_4_, *c*_5_, and *c*_6_ in every environment – *c*_1_ is universally better than these cultivars, and ranking any of *c*_3_, *c*_4_, *c*_5_, or *c*_6_ above *c*_1_ is a violation of Pareto efficiency.

**TABLE 1.**
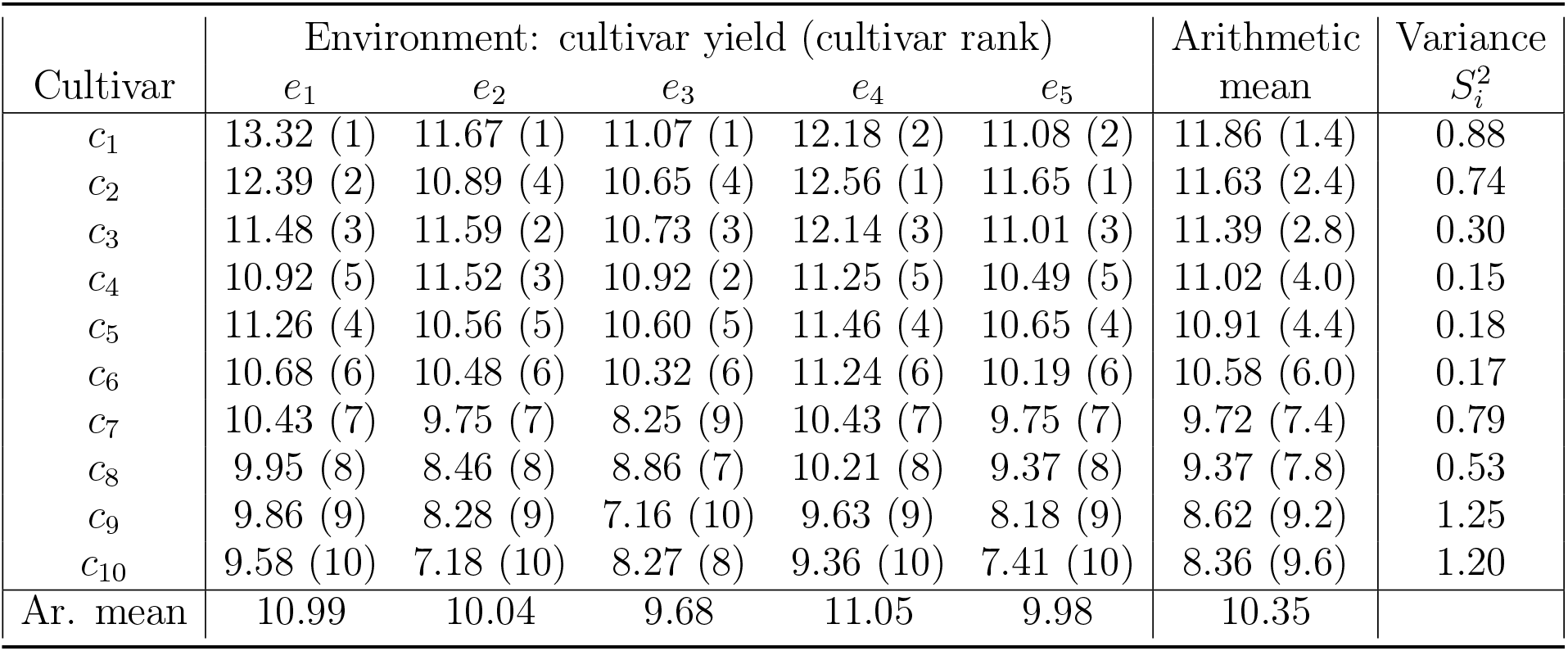
The yield profile of a multi-environment trial.

**TABLE 2.** Cultivar rankings under yield stability metrics.

| | $S^{(1)}$ | $S^{(2)}$ | $S^{(3)}$ | $S^{(6)}$ | $NP^{(1)}$ | $NP^{(2)}$ | $NP^{(3)}$ | $NP^{(4)}$ | $W_i^2$ | $\sigma_i^2$ | $\theta_i$ | $\theta'_i$ | $S_i^2$ | $CV_i$ | AR |
| --- | --- | --- | --- | --- | --- | --- | --- | --- | --- | --- | --- | --- | --- | --- | --- |
| $c_1$ | 5.5 | 5.5 | 3 | 3 | 6.5 | 3 | 4 | 4 | 7 | 7 | 7 | 7 | 8 | 7 | 5 |
| $c_2$ | 10 | 10 | 8 | 6 | 5 | 4 | 3 | 3 | 3 | 3 | 3 | 3 | 6 | 5 | 4 |
| $c_3$ | **3** | **3** | **2** | **2** | **3** | **1.5** | **2** | **2** | 5 | 5 | 5 | 5 | **4** | **4** | **3** |
| $c_4$ | 9 | 9 | 9 | 7 | 9.5 | 7 | 6 | 6 | 9 | 9 | 9 | 9 | **1** | **1** | 7 |
| $c_5$ | 5.5 | 5.5 | 4 | 4 | **1** | 1.5 | **1** | **1** | **1** | **1** | **1** | **1** | 3 | 2 | **1** |
| $c_6$ | **1** | **1** | **1** | **1** | 3 | 5 | 5 | 5 | 2 | 2 | 2 | 2 | 2 | 3 | 2 |
| $c_7$ | 7.5 | 7.5 | 7 | 8.5 | 6.5 | 8 | 8 | 8 | 6 | 6 | 6 | 6 | 7 | 8 | 8 |
| $c_8$ | 3 | 3 | 5 | 5 | 3 | 6 | 7 | 7 | 4 | 4 | 4 | 4 | 5 | 6 | 6 |
| $c_9$ | 3 | 3 | 6 | 8.5 | 8 | 9 | 9 | 9 | 8 | 8 | 8 | 8 | 10 | 9 | 9 |
| $c_{10}$ | 7.5 | 7.5 | 10 | 10 | 9.5 | 10 | 10 | 10 | 10 | 10 | 10 | 10 | 9 | 10 | 10 |

*Notes:* The cultivar marked with 1 is the most stable under a given metric. The calculations are provided in Appendix C. The bold entries highlight that the frontier obtained from mean yield and a stability statistic can include cultivars that are Pareto-dominated in the original yield profile. Hence, summarizing performance by mean yield and stability can violate Pareto efficiency.

Pareto efficiency alone is not sufficient. If it were, this problem would have an easy solution – first throw away the Pareto inefficient cultivars, then apply the stability measures to what remains. But if we take this approach, we will violate some fundamental consistency requirements. For the sake of argument, suppose our strategy is to drop the obviously bad cultivars to maintain Pareto efficiency, and then rank the remaining cultivars based on variance of yield. If we throw away the cultivars obviously inferior to *c*_1_, we are left with *c*_1_ and *c*_2_. Cultivar *c*_2_ has a lower variance than *c*_1_, hence *c*_2_ is the cultivar that our rule recommends. Now suppose that the yield of *c*_1_ in environment *e*_1_ was a bit worse – not 13.32, but only 12.32. In this case, *c*_1_ would have a lower variance (0.35) than *c*_2_, and therefore *c*_1_ would be selected. Paradoxically, in the eyes of this two-step ranking procedure, a worse yield for *c*_1_ makes it a better cultivar.

If the yield of *c*_1_ in environment *e*_1_ were even lower, only 10.32, then the outcome becomes even stranger. Cultivar *c*_1_ is no longer obviously better than *c*_3_ and *c*_4_, since *c*_3_ and *c*_4_ have a higher yield in environment *e*_1_. Considering all four cultivars, *c*_4_ has the lowest variance and is chosen by our rule. However, observe that the only difference between the two multi-environment trials is the yield of *c*_1_. Why should the yield of *c*_1_ determine which of *c*_2_ or *c*_4_ is chosen?

A ranking rule with such paradoxical properties is difficult to recommend. The relative ranking of two cultivars should depend on their own yields, not on the presence, absence, or yield of unrelated cultivars; increasing the yield of a cultivar should not harm its ranking; environments that provide no discriminatory information should not affect the final order; and changes in measurement units should not change cultivar rankings (Luce and Raiffa, 1957; Knight, 1970; Sen, 1979; Roberts, 1980; Mead et al., 1986). These requirements are not merely abstract mathematical preferences. They correspond to practical expectations that breeders, agronomists, seed companies, and farmers implicitly place on cultivar recommendation procedures.

The axiomatic approach allows us to evaluate ranking rules according to these consistency requirements. The example above shows that several commonly used stability-based ranking procedures can fail basic axioms, including weak Pareto efficiency or independence of irrelevant alternatives, but in this paper we show something stronger: *every* conceivable ranking procedure, with the exception of a single coherent class, fails to satisfy a minimal combination of desirable properties. The exception is the family of power means. This family includes the arithmetic mean, geometric mean, harmonic mean, minimum rule, and maximum rule as members or limiting cases. The parameter of the power mean controls the degree to which the rule rewards high yields or penalises low yields, which allows the breeder to choose a ranking procedure that balances high average yield with the breeder’s appetite for risk.

The geometric mean occupies a central position in this framework. It preserves the intuitive appeal of an average while penalising low yields more strongly than the arithmetic mean (Grabisch et al., 2011). A cultivar with catastrophic yield in one environment cannot fully compensate for that failure with high yield elsewhere. This behaviour reflects the risk-aversion of farmers, for whom unusually low yields may be more damaging than unusually high yields are beneficial (Mishra and Sandretto, 2002; Cernay et al., 2015; Macholdt and Honermeier, 2016; Duvallet et al., 2021; von Gehren et al., 2023). In addition, the geometric mean is naturally compatible with multiplication normalisation of environmental scale, in which cultivar yield is evaluated relative to the productivity level of each environment.

This paper contributes to the cultivar-selection literature in three ways. First, it reformulates the simultaneous evaluation of yield potential and yield stability as the selection of a ranking rule rather than the selection of a stability statistic. Second, it adapts an axiomatic approach from social choice theory and operations research to evaluate the logical consistency of cultivar-ranking rules (Luce and Raiffa, 1957; Arrow, 1963; Sen, 1979; Roberts, 1980; Moulin, 1988; Chebotarev and Shamis, 1998; Finan and Hurley, 2002; Bossert and Weymark, 2004; Fortemps and Pirlot, 2004). Third, it provides a principled justification for the geometric mean, and more generally for lower-order power means, as ranking rules that combine productivity and stability while satisfying desirable axiomatic properties.

The objective is therefore not to replace established statistical methods for genotype-by-environment interaction. Eberhart–Russell regression, Finlay–Wilkinson regression, Shukla stability variance, AMMI, SREG/GGE biplots, shifted multiplicative models, mixed models, factor-analytic models, and genomic prediction models remain essential for estimating cultivar performance, characterising adaptation patterns, identifying mega-environments, modelling crossover interaction, and predicting unobserved genotype-by-environment combinations. The present paper begins where these methods leave off: once cultivar yield across environments has been observed, adjusted, estimated, or predicted, how should cultivars be ranked for recommendation?

The remainder of the paper is organized as follows. In Section 2.1 we introduce the formal model for complete multi-environment trials with fixed sets of cultivars and environments, define ranking rules, and in Section 2.2 evaluate several desirable axioms. In Sections 2.3 and 2.4 we extend the axiomatic approach to trials with variable sets of cultivars and environments, and in Sections 2.5 and 2.6 consider normalised and incomplete data, showing how different normalisation methods interact with the axioms and why the geometric mean is singled out under multiplication normalisation. In Section 3 we discuss the implications for cultivar recommendation, yield stability analysis, and the practical use of power means in plant breeding. Appendix A contains related results obtained from axioms that are less relevant in a cultivar selection context; the proofs of the theorems can be found in Appendix B.

## 2. Methods

### 2.1. Ranking methods for fixed size trials

We want to formalise the process of ranking cultivars on the basis of a multi-environment trial in a mathematically precise way. We will use *c*_*i*_ to denote the *i*th cultivar and *e*_*j*_ the *j*th environment. The yield of *c*_*i*_ in *e*_*j*_ is denoted by *y*_*ij*_.

A **yield profile Y** is a matrix of quantities *y*_*ij*_ indexed by a finite set of cultivars *C* ⊂ **N** and a finite set of environments *E* ⊂ **N, Y** = {*y*_*ij*_}_*i*∈*C, j*∈*E*_. A yield profile could be thought of as the yield data of a concrete multi-environment trial.

In this subsection we assume that the set of cultivars and environments is fixed in advance and cannot vary between yield profiles. We will consider the case of variable size yield profiles in Subsections 2.3-2.4. Another consequence of our definition is that the trial is complete – every cultivar has a recorded yield in every environment. We will touch a bit upon incomplete trials in Subsections 2.5-2.6.

A **ranking rule** is a function that, given a yield profile **Y** on (*C, E*), produces a ranking (complete and transitive order) of cultivars from *C*. The outcome of the ranking rule is called the final ranking and may contain ties between cultivars.

We impose the following condition on the space of allowable ranking rules in this paper: if cultivars *c*_*i*_ and *c*_*k*_ have the same yield in every single environment, *y*_*ij*_ = *y*_*kj*_ for all *j*, then the ranking rule must rank *c*_*i*_ and *c*_*k*_ equal to each other. While we could come up with mathematically well defined procedures that violate this criterion, such procedures clearly do not reflect the intention that we want the ranks of the cultivars to somehow reflect their performance.

The usual yield stability measures used in the literature, both parametric and non-parametric, fall under this definition of a ranking rule, as does just about any conceivable procedure. The only real restriction is that our definition of a ranking rule is deterministic – we do not allow tossing a coin to help us determine a ranking (Tohidi and Olafsson, 2025). We will give a couple of examples.

The simplest measure of yield stability is variance, 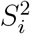. For the yield profile in Table 1, we give the variance scores in a separate column, so the the best cultivar is *c*_4_, followed by *c*_6_, *c*_5_, *c*_3_, *c*_8_, *c*_2_, *c*_7_, *c*_1_, *c*_10_, and *c*_9_.

As we have seen in the introduction, yield stability measures such as this run the risk of choosing a “universally bad” cultivar if that cultivar happens to be bad in a stable way. Let us fix this with a refinement of the 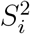 ranking rule.

We say that *c*_*i*_ **Pareto-dominates** *c*_*k*_ and *c*_*k*_ is **Pareto-dominated** by *c*_*i*_, if *y*_*ij*_ ≥ *y*_*kj*_ for each *e*_*j*_, and there exists at least one *e*_*l*_ such that *y*_*il*_ *> y*_*kl*_.

The ranking rule 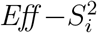 first sets all the Pareto-dominated cultivars aside, and ranks all the undominated cultivars in terms of their variance. Then it takes the cultivars that are left, sets aside those that are dominated within the group that is left, and ranks the undominated in terms of their variance. This process continues until all cultivars are ranked.

In the yield profile in Table 1, the undominated cultivars are *c*_1_ and *c*_2_. First we rank these, and get *c*_2_, *c*_1_. Looking at the remaining cultivars, *c*_3_ and *c*_4_ are undominated. If we rank them in terms of their variance we get *c*_4_, *c*_3_. Continuing this process, the final ranking is *c*_2_, *c*_1_, *c*_4_, *c*_3_, *c*_5_, *c*_6_, *c*_8_, *c*_7_, *c*_10_, *c*_9_.

The **TOP ranking rule** is an interesting example of a yield stability metric in the literature that does not explicitly use any statistical measures of the yield data. Instead it ranks a cultivar according to the percentage of environments for which this cultivar is within the top third of cultivars (Fox et al., 1990). On the same yield profile, this rule would rank *c*_1_ and *c*_3_ tied first, then *c*_2_, then *c*_4_, then all the others tied last.

To conclude this section, let us introduce a family of purely mathematical ranking rules that generalise the concept of the mean.

Let *w*_1_, …, *w*_|*E*|_ be the **weights** of the environments – a sequence of non-negative numbers such that :

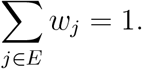

The **weighted power mean** (weighted root-mean-power) of the yield of *c*_*i*_ is defined with respect to a parameter −∞ *< α <* ∞ as (Grabisch et al., 2011):

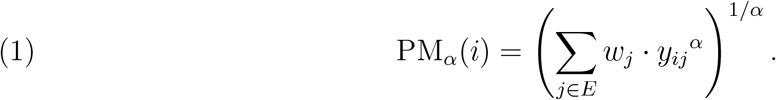

If all the weights are equal, this is called the (unweighted) **power mean**.

Taken as a ranking rule, the weighted power mean ranks cultivars with respected to the weighted power mean of their yields. Observe that with *α* = 1 this is equivalent to ranking cultivars in terms of their mean yield, the **weighted arithmetic mean**:

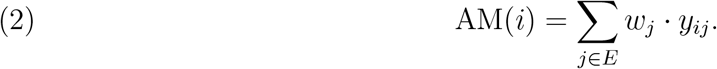

This is a very natural ranking rule, but generally seen to be inappropriate for ranking cultivars because unstable cultivars with potentially catastrophic yields in some circumstances can nevertheless have a high arithmetic mean (Mohammadi and Amri, 2008; Reckling et al., 2021).

As *α* → 0, the formula above can be simplified to the more familiar **weighted geometric mean**, which ranks cultivars in accordance with the weighted product of their yields:

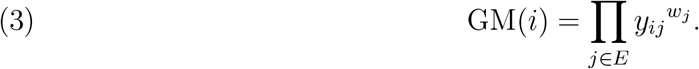

The geometric mean was motivated in the plant breeding literature by Mohammadi and Amri (2008). Note that the geometric mean punishes cultivars with occasional bad performance more than the arithmetic mean – indeed, a yield of 0 in a single environment would result in a geometric mean of 0, while the arithmetic mean could be compensated by good yields in other environments. This is the intuitive explanation of *α*. When *α* is small, we disproportionately punish cultivars for poor performance in some environments. When *α* is large, we disproportionately reward cultivars for good performance in some environments. Table 1 has a column for the arithmetic mean (*α* = 1). This rule would rank the cultivars *c*_1_, *c*_2_, *c*_3_, *c*_4_, *c*_5_, *c*_6_, *c*_7_, *c*_8_, *c*_9_, *c*_10_.

To generalise this idea of punishing bad performance or rewarding good performance, we also consider the **maximum** and **minimum** ranking rules, which rank cultivars by their single best/worst yield.

### 2.2. Axioms and theorems for fixed size trials

In the previous section we formalised three yield stability measures from the literature as ranking rules (the TOP rule, 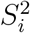, and 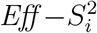), and defined an infinite family of ranking rules, representing different ways of extending the concept of the mean. As the reader can appreciate, the only limit in defining new rules is our imagination. What we would like is a way to distinguish between rules which have desirable properties and those that do not.

We will start by revisiting the notion of the “universally bad” cultivars from the introduction.

A ranking rule is **weakly Pareto efficient** if for each yield profile where *y*_*ij*_ ≥ *y*_*kj*_ for each *j*, it ranks *c*_*i*_ higher than or equal to *c*_*k*_. A ranking rule is **Pareto efficient** if it is weakly Pareto efficient and for each yield profile where *y*_*ij*_ *> y*_*kj*_ for each *e*_*j*_, it ranks *c*_*i*_ higher than (and not equal to) *c*_*k*_ (Chebotarev and Shamis, 1998). In the literature on plant breeding, Pareto efficiency was motivated by Mead et al. (1986).

The difference between weakly and properly Pareto efficient rules is decisiveness – the rule which always ranks all cultivars as equal is weakly Pareto efficient, but it is not a very useful rule.

We have already seen that 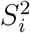 is not even weakly Pareto efficient. In the appendix we show that the same is true of almost every yield stability measure used in the literature. The TOP rule is weakly Pareto efficient, but not Pareto efficient – in Table 1, cultivar *c*_1_ has a better yield in every environment than *c*_3_, but TOP ranks them tied for first place. 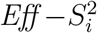 is Pareto-efficient, as is any rule in the family of weighted power means.

The other notion we saw in the introduction was **Independence of Irrelevant Alternatives** (IIA). A ranking rule satisfies Independence of Irrelevant Alternatives if changing the yield of a cultivar in a yield profile leaves the relative ranking of the other alternatives unchanged (Sen, 1979, p. 129, first published in 1970). The idea here is that the ranking of cultivars *a* and *b* should depend purely on the relative merits of *a* and *b*, and not whether we consider cultivar *c* as well.

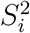 and the power means satisfy IIA. We have already seen that 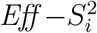 does not. Interestingly, neither does TOP – if we lower the yield of *c*_3_ in *e*_1_ to 11.2, then *c*_5_ will take *c*_3_’s place in the top third of cultivars in *e*_1_, and will be ranked above *c*_6_, *c*_7_, *c*_8_, *c*_9_, *c*_10_.

Independence of Irrelevant Alternatives is a cultivar-level notion of independence: irrelevant cultivars should not affect the ranking. We can also consider environment-level independence – if an environment tells us nothing important, then it should not affect the outcome of the ranking rule. The most obvious example of an informationally useless environment is some *e*_*j*_ such that *y*_*ij*_ = *y*_*kj*_ for all *i, k* – every cultivar has the same yield. For example one could think of *e*_*j*_ being an arctic waste where *y*_*ij*_ = *y*_*kj*_ = 0. We call such an environment **indifferent**. A ranking rule satisfies **separability** if the ranking of the cultivars is unchanged if we change the yield (of all cultivars) on an indifferent environment (Roberts, 1980).

The geometric mean, without further refinements, fails separability. In the case of arctic wasteland above, all cultivars would be tied with a geometric mean of zero. However, if we increase the yield in this environment to a non-zero value, we could get a very different ordering of the cultivars. It is possible to recover separability for the geometric mean by introducing an appropriate tie-breaking mechanism in the case of zero yields; we will return to the idea of such a refinement shortly.

The last property we want to look at in this section is **multiplication invariance**. A ranking rule satisfies multiplication invariance if multiplying the yield of every single cultivar in every single environment by the same constant *x* will leave the ranking of the cultivars unchanged (Roberts, 1980). One way of interpreting multiplication invariance is unit invariance: if we change the measurement units from pounds to kilograms, it is the equivalent of multiplying all yields by approximately 0.45. We would expect that the ranking of the cultivars would be unchanged by a change of units. All the measures we found in the literature satisfy multiplication invariance.

We are almost ready to see where these properties lead us, but first we want to formalise the notion of two ranking rules being almost the same, but one being more particular in how it orders cultivars that are hard to distinguish from each other.

A ranking rule *R* is a **refinement** of *Q* if whenever *Q* ranks *c*_*i*_ above *c*_*k*_, then *R* ranks *c*_*i*_ above *c*_*k*_ as well. However, if *c*_*i*_ is ranked equal to *c*_*k*_ according to *Q*, then *R* may order *c*_*i*_ and *c*_*k*_ any way it likes – either *c*_*i*_ could be ranked higher, or *c*_*k*_ could be ranked higher, or *R* could agree with *Q* and rank them equal to each other. Thus any ranking rule is a refinement of itself.

#### Theorem 1.

(Roberts 1980, p. 432). Consider a domain with positive yields, complete yield profiles, and fixed sets of cultivars and environments with at least three cultivars. If a ranking rule satisfies multiplication invariance, Pareto efficiency, independence of irrelevant alternatives, and separability, then it is a refinement of either the minimum rule, the maximum rule, or the weighted power mean.

#### Breeding interpretation of Theorem 1

If a cultivar ranking rule is not affected by yield units, respects uniformly superior cultivars, gives independent pairwise comparisons, and ignores informationally useless environments, then it must belong to the weighted power mean family or its limiting cases.

Theorems such as the above demonstrate the power of the axiomatic approach. There is an infinity of feasible ranking rules out there, but we do not need to consider every single one. By imposing a handful of desirable properties, we are left with a single, intuitive family, and two edge cases.

As the reader can gather, if we impose more axiomatic properties, we will end up with a smaller set of admissible rules. The property of **reversibility** captures the notion that exceptionally bad yields are potentially catastrophic events and should not be offset by good yields in other environments. A ranking rule satisfies reversibility if for any yield profile with positive yields, cultivar *c*_*i*_ and environment *e*_*j*_, it is possible to decrease *y*_*ij*_ to the point where *c*_*i*_ will be ranked last in the final ranking, for example by setting *y*_*ij*_ very close to zero.^1^ If we consider this to be a sensible property, we can get rid of the maximum rule and the convex means.

##### Theorem 2.

Consider a domain with positive yields, complete yield profiles, and fixed sets of cultivars and environments with at least three cultivars. If a ranking rule satisfies multiplication invariance, Pareto efficiency, independence of irrelevant alternatives, separability, and reversibility, then it is a refinement of the minimum rule or the weighted power mean with parameter *α* ≤ 0 and positive weights.

#### Breeding interpretation of Theorem 2

If a cultivar ranking rule is not affected by yield units, respects uniformly superior cultivars, gives independent pairwise comparisons, ignores informationally useless environments, and protects against cultivars with occasional severe failure, then it must belong to the minimum rule or the weighted power mean family with parameter *α* ≤ 0, including geometric and harmonic means.

### 2.3. Ranking methods for variable size trials

In this section, we generalise the model to take into account the fact that different trials can cover different cultivars and environments, and different numbers of cultivars and environments. In our mathematical language, each yield profile **Y** can be indexed by a different set of environments *E* ⊂ **N** and cultivars *C* ⊂ **N**, and a ranking rule must produce a ranking for any such yield profile. To see why this makes a difference, let us consider the form of the weighted power mean in this setting.

Let 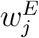 be a set of non-negative numbers such that:

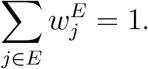

The weighted power mean is defined with respect to a parameter −∞ *< α*_*E*_ *<* ∞ as:

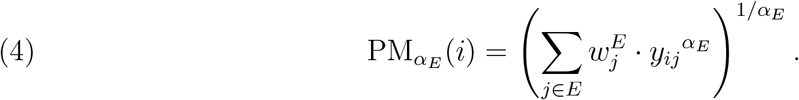

The weighted power mean for *α*_*E*_ = 0 is the **weighted geometric mean** defined as:

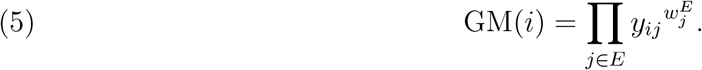

Compared to the definition in (1), there are two differences:

1. The set of weights 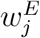 is defined with respect to a given set of environments. For example, we might use the weights 0.4 and 0.6 if the trial takes place in a dry and a wet location, 0.3 and 0.7 if it takes place in a cold and a hot location, and 0.2, 0.3, 0.5 if the environments are low, medium, and high levels of pesticide.
2. The parameter *α*_*E*_ is also a function of *E* and not a constant. So we could use the arithmetic mean (*α* = 1) in the case of a dry and a wet location, the geometric mean (*α* = 0) in the case of a cold and a hot location, the harmonic mean (*α* = −1) in the case of different levels of pesticide, and whatever else we can come up with. We are not necessarily claiming there are sensible reasons for doing so, but the model is flexible enough to allow it.

### 2.4. Axioms and theorems for variable size trials

In this setting we can define Independence of Irrelevant Alternatives, which we call **strict Independence of Irrelevant Alternatives** (SIIA), in a more intuitive way. A ranking rule satisfies strict independence of irrelevant alternatives if the removal of a cultivar from a yield profile leaves the relative ranking of the other cultivars unchanged (Luce and Raiffa, 1957, p. 343). In the literature on plant breeding, the importance of SIIA was highlighted by Knight (1970) and Mead et al. (1986).

#### Theorem 3.

Consider a domain with positive yields, complete yield profiles, and *variable* sets of cultivars and environments. If a ranking rule satisfies multiplication invariance, Pareto efficiency, *strict* independence of irrelevant alternatives, and separability, then:

1. for each set of environments, it is a refinement of the minimum rule, maximum rule, or weighted power mean.
2. if it also satisfies reversibility, then for each set of environments *E*, it is a refinement of the minimum rule or the weighted power mean with parameter *α*_*E*_ ≤ 0 and positive weights 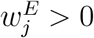.

The practical interpretation of Theorem 3 is identical to Theorems 1 and 2.

### 2.5. Normalisation methods

In practice, the yield profile is often incomplete. A cultivar *c*_*i*_ may not have been measured in environment *e*_*j*_ and the yield *y*_*ij*_ is missing, or perhaps the yield was measured, but there is an obvious error in the data, so it was decided not to use the result (Boehm Jr et al., 2019; Laidig et al., 2021; Reckling et al., 2021; Wang et al., 2021; Abdala et al., 2024; Ramakers et al., 2025).

Most of the measures in the literature can be adapted to this situation easily enough: we can compute the average yield or variance for cultivars *c*_*i*_ and *c*_*k*_ separately, even if the cultivars were grown in different environments. However, the comparison in this case may not be meaningful: if cultivar *c*_*i*_ was evaluated in well-irrigated environments and cultivar *c*_*k*_ was not, then the average yield of *c*_*i*_ would appear to be higher, but that doesn’t necessarily mean *c*_*k*_ wouldn’t do as well in the same conditions. In other applications, the usual way to correct for this is to normalise the initial data (Schenkerman, 1994; Munda, 2008; Tofallis, 2012). We will consider the following popular methods to normalise the yield:

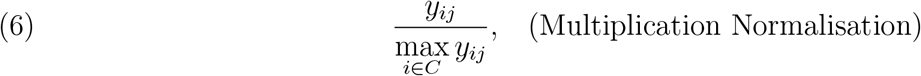

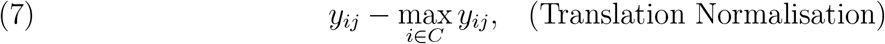

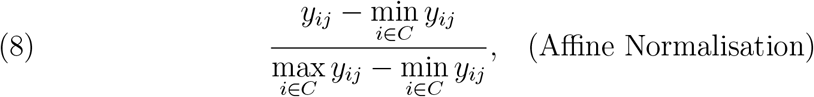

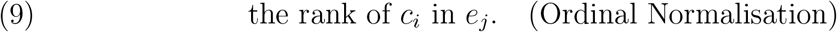

In the literature on plant breeding, translation normalisation was used by Lin and Binns (1988) and ordinal normalisation is very common (Mohammadi and Amri, 2008; Pour-Aboughadareh et al., 2022).

### 2.6. Axioms and theorems for normalisation methods

We say that a ranking rule **admits normalisation** N if the outcome is the same for all yield profiles that coincide after normalisation N. In other words, rule *R* admits normalisation N if *R*(**Y**) = *R*(**Y**^*′*^) for all profiles **Y** and **Y**^*′*^ such that N(**Y**) = N(**Y**^*′*^).

The following theorem shows that if we are interested in ranking rules that admit multiplication or translation normalisation, we are essentially limited to a refinement of the geometric or arithmetic mean.

#### Theorem 4.

Consider a domain with positive yields, complete yield profiles, and *fixed* sets of cultivars and environments with at least three cultivars. If a ranking rule satisfies Pareto efficiency and independence of irrelevant alternatives, then:

1. if it also satisfies separability and admits multiplication normalisation (6), then it is a refinement of the weighted geometric mean (follows from a result on p. 432 in Roberts 1980).
2. if it also satisfies separability, reversibility, and admits multiplication normalisation (6), then it is a refinement of the weighted geometric mean with positive weights.
3. if it also satisfies multiplication invariance and admits translation normalisation (7), then it is a refinement of the weighted arithmetic mean (follows from Roberts 1980, theorem 2).

In the case of *variable* sets of cultivars and environments, if a ranking rule satisfies Pareto efficiency and *strict* independence of irrelevant alternatives, then:

(4) if it also satisfies separability and admits multiplication normalisation (6), then, for each set of environments, it is a refinement of the weighted geometric mean.
(5) if it also satisfies separability, reversibility, and admits multiplication normalisation (6), then, for each set of environments *E*, it is a refinement of the weighted geometric mean with positive weights 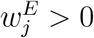.
(6) if it also satisfies multiplication invariance and admits translation normalisation (7), then, for each set of environments, it is a refinement of the weighted arithmetic mean.

#### Breeding interpretation of Theorem 4

Multiplication normalisation singles out the weighted geometric mean. Translation normalisation singles out the weighted arithmetic mean; this is undesirable in multi-environment trials because the arithmetic mean allows poor yields in some environments to be easily offset by good yields in others.

This result, while restrictive, nevertheless has a positive message: if we are interested in using multiplication normalisation, then there does exist a ranking rule that satisfies us – the geometric mean. Our next two theorems are a lot more dire, and will show that many combinations of normalisation methods and desirable properties are simply incompatible.

One of these properties is the idea that all environments should be informationally useful, and we should not allow any one environment to dictate the outcome. For a ranking rule and a set of at least two cultivars and at least two environments, an environment is called a **weak dictator** if, whenever the yield of cultivar *c*_*i*_ is higher than the yield of cultivar *c*_*k*_ in this environment, then *c*_*i*_ will be ranked higher than *c*_*k*_ in the final ranking. In other words, the rule is a refinement of the weighted power mean where only one environment (the weak dictator) has positive weight. The presence of a weak dictator is a very negative feature, because it negates the whole point of a multi-environment trial: if we only care about one environment, why waste the time and resources to conduct the trial in others?

##### Theorem 5.

Consider a domain with positive yields, complete yield profiles, and *fixed* sets of cultivars and environments with at least three cultivars. If a ranking rule satisfies Pareto efficiency, independence of irrelevant alternatives, and admits affine (8) or ordinal (9) normalisation, then there exists a weak dictator (follows from Arrow 1963, p. 97, theorem 8*2 in Sen 1979, first published in 1970, and theorem 3 in Roberts 1980).

In the case of *variable* sets of cultivars and environments, if a ranking rule satisfies Pareto efficiency, *strict* independence of irrelevant alternatives, and admits affine (8) or ordinal (9) normalisation, then, for each set of environments, there exists a weak dictator.

#### Breeding interpretation of Theorem 5

Affine or ordinal normalisation methods cause a single environment to dictate the ranking. This is undesirable in multi-environment trials because cultivar recommendation should reflect performance across environments.

The second property is **monotonicity**, which captures the idea that increasing the yield of a cultivar should not harm its ranking. Formally, we say that a ranking rule is monotonic if whenever *c*_*i*_ is ranked above *c*_*k*_ and equal to *c*_*l*_ by the ranking rule, and we increase the yield of *c*_*i*_ in *e*_*j*_, then *c*_*i*_ is still ranked above *c*_*k*_, and is ranked equal to or higher than *c*_*l*_ (Luce and Raiffa, 1957, p. 343).

Monotonicity is a strengthening of weak Pareto efficiency: every monotonic rule is also (at least) weakly Pareto efficient, but there exist weakly Pareto efficient rules that are not monotonic. As a result monotonicity is a harder property to satisfy – we have seen that we can obtain Pareto efficiency by essentially getting rid of the inefficient cultivars, as with the rule 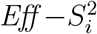, but such multi-step procedures are typically not monotonic, as indeed 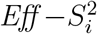 is not. The family of weighted power means, however, is monotonic, so monotonicity is consistent with all the preceding theorems.

##### Theorem 6.

Consider a domain with positive yields, complete yield profiles, and *variable* sets of cultivars and environments. Then:

1. no ranking rule that admits translation normalisation (7) satisfies reversibility and monotonicity.
2. no ranking rule that admits affine (8) or ordinal (9) normalisation satisfies reversibility.

#### Breeding interpretation of Theorem 6

Translation, affine, or ordinal normalisation methods conflict with protection against catastrophic yield failures.

Our final result deals with yield profiles that have some data missing, and it shows that in this context no reasonable normalisation method is consistent with independence of irrelevant alternatives.

We call an environment *e*_*j*_ **almost indifferent** if *y*_*ij*_ = *y*_*kj*_ for all *i, k*, where the yield was measured and used – every cultivar has the same yield. A normalisation method is said to be **neutral** if any two yield profiles, where missed data are the same and all environments are almost indifferent, coincide after normalisation.

We say that a ranking rule is **imposed** if the relative ranking of two cultivars, whose yield was measured and used in disjoint sets of environments, does not depend on the numbers in the yield profile.

##### Theorem 7.

Consider a domain where yield profiles are allowed to be incomplete. If a ranking rule satisfies the independence of irrelevant alternatives and admits some neutral normalisation method, then it is imposed.

#### Breeding interpretation of Theorem 7

Whenever statistical models produce reliable adjusted, estimated, or predicted yields, these yields should be used as input to ranking rules.

## 3. Results and Discussion

The objective of this paper was to reconsider cultivar recommendation in multi-environment trials as a ranking problem. The outcome of the trial is that a cultivar is associated with a vector of yields across environments rather than by a single value. However, breeding decisions ultimately require an ordering of cultivars for advancement, release, promotion, or rejection. This final step is often treated as a natural consequence of mean yield, a stability statistic, or a graphical genotype-by-environment analysis. We argue that it should instead be treated as a distinct decision problem requiring explicit formal justification. The approach we take is that of axiomatic ranking theory: we formally define properties that a “reasonable” ranking rule should satisfy, and prove theorems to show how the space of admissible ranking rules is restricted depending on which properties we impose.

A central message of the paper is that yield stability analysis and cultivar ranking are not identical problems. Classical stability measures, including variance-based statistics, regression-based parameters, and non-parametric indices, are useful for describing stability of yield across environments. Similarly, AMMI, SREG/GGE biplots, linear-bilinear models, shifted multiplicative models, mixed models, factor-analytic models, genomic prediction models, and high-throughput phenotyping provide powerful tools for estimating, decomposing, visualising, or predicting genotype-by-environment interaction. These methods help breeders understand adaptation patterns, detect crossover interaction, identify mega-environments, estimate cultivar effects, and predict unobserved genotype-by-environment combinations. However, they do not automatically define how the resulting yield profiles should be aggregated into a final recommendation ranking.

The illustrative multi-environment trial in Table 1 makes this distinction concrete. In the example, cultivar *c*_1_ has the highest yield in three environments and the second-highest yield in the remaining two environments. More importantly, *c*_1_ outperforms cultivars *c*_3_ through to *c*_10_ in *every single* environment. By any reasonable standard, it is the better cultivar. However, in Table 2 we demonstrate that many commonly used stability metrics identify these inferior cultivars as more stable. These cultivars have lower variability, but their apparent stability is partly a consequence of consistently lower yield. Thus, the example illustrates the central problem addressed in this paper: a method can correctly describe one aspect of stability and still be inappropriate as a final recommendation rule if it ignores that some cultivars have universally higher yield. A defensible ranking rule must combine stability and reliability while respecting minimal efficiency requirements such as this.

This property – that a universally higher yielding cultivar should be ranked at least as high as a universally lower yielding cultivar – is known as weak Pareto efficiency, and it is a minimal principle for cultivar recommendation. A cultivar that is uniformly dominated should not be recommended merely because its yield is stable.

The concept of Pareto efficiency has already been widely used in plant breeding, if never as a principle for cultivar ranking. It is implicit when balancing between the probability of success, cost, and time in in gene stacking (Xu et al., 2011) and trait introgression problems (Cameron et al., 2017); balancing between gain, risk, and inbreeding in genomic mating design (Akdemir and Sánchez, 2016; Hunter and McClosky, 2016); balancing the desirable traits of cultivars, e.g. yield, height, and protein content (Akdemir et al., 2019). Closer to the current paper, Pareto efficiency has been useful in selecting discriminating test locations in multi-environment trials – the tradeoff is between the traditional measure of precision and the relative discriminative value of a location (Kiaghadi and Olafsson, 2025).

Pareto efficiency is a very minimal requirement and satisfying Pareto efficiency does not in itself mean that the ranking rule is a good one. However, the remarkable fact is that Pareto efficiency combined with just a handful of other minimal requirements will collapse the space of admissible rules to a single family. If we impose multiplication invariance – the requirement that the ranking should not change if we scale all yields by a common constant, independence of irrelevant alternatives – that the relative ranking of two cultivars should not depend on the yield of a third cultivar, and separability – the ranking should not depend on informationally useless environments, then we are left with the family of power means and their limiting cases (Theorem 1).

The power means are ubiquitous in mathematics, and include the well known arithmetic and geometric means as special cases. That said, few would argue that the arithmetic mean is an appropriate rule to rank cultivars – the arithmetic mean allows poor yields in some environments to be easily offset by good yields in others, but farmers typically face asymmetric risks from poor harvests, which is why such a focus is placed on the stability of yield in the literature. In our framework, we capture this with the property of reversibility – the property that a catastrophic yield in one environment cannot be offset by high yields elsewhere. If we add reversibility to the properties satisfied by the ranking rule, we are left with power means with parameter *α* ≤ 0 (Theorem 2); the geometric mean is admitted by this theorem, while the arithmetic mean is not.

When faced with incomplete data, it makes sense to normalise the yield profiles to account for the fact that some cultivars may have been tested in high yield environments, e.g. with high fertiliser treatment, while others were not. In the context of normalisation a natural property to impose is that the ranking of the cultivars before and after yield normalisation is unchanged. With this requirement, we are essentially left with the geometric mean as the uniquely suitable ranking rule (Theorems 4, 5, 6). We summarise our results in Table 3, and in Table 4 we show which combination of properties pins down which ranking rule.

**TABLE 3.**
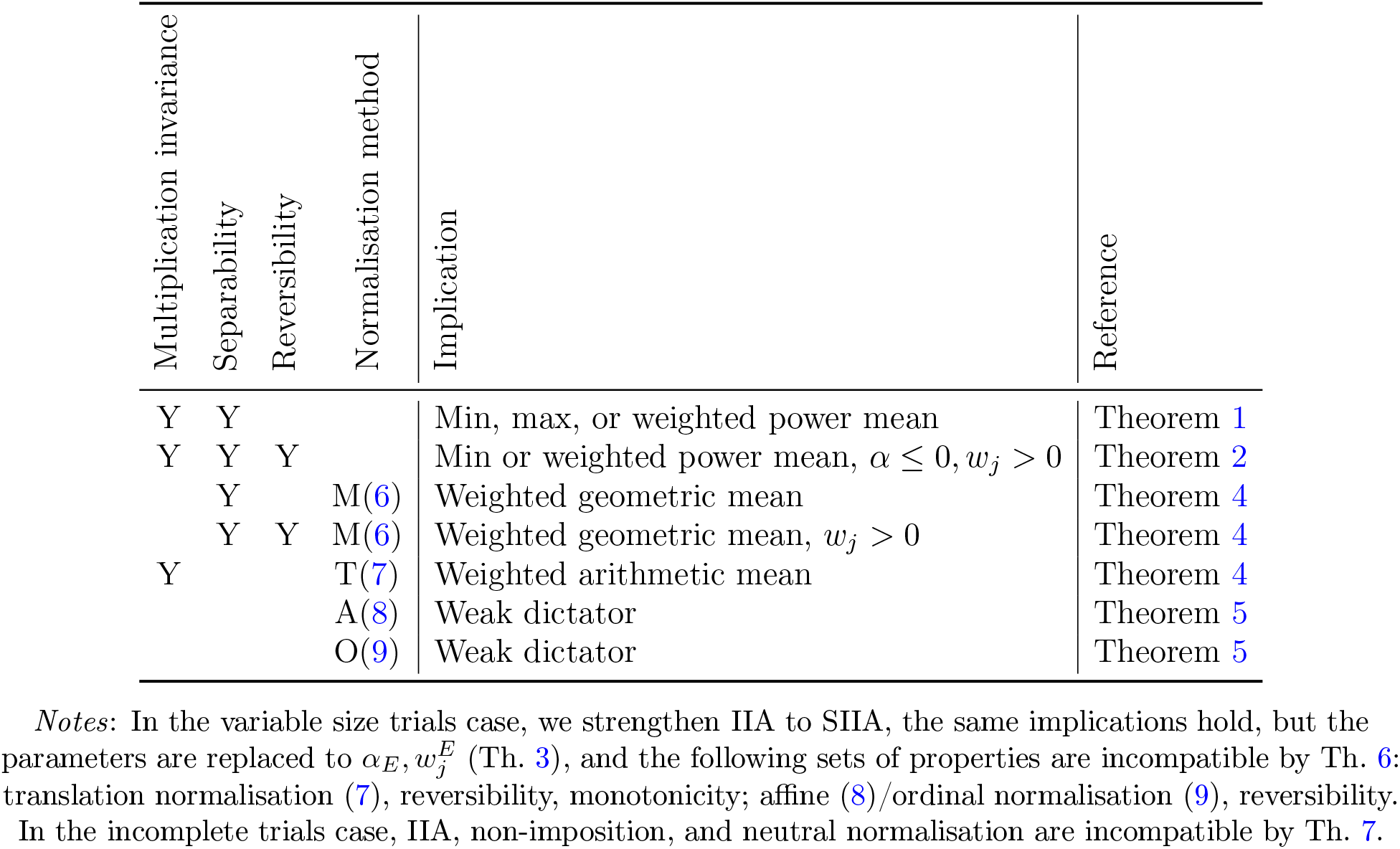
Implications of IIA, Pareto efficiency, and other axioms: fixed size trials.

**TABLE 4.** Comparison of ranking rules: complete yield profiles.

| Multiplication invariance | Separability | Weak Pareto efficiency | Monotonicity | Pareto efficiency | IIA, SIIA | Reversibility | Admits normalisation | Ranking rule |
| --- | --- | --- | --- | --- | --- | --- | --- | --- |
| Y | Y | Y | Y | Y | Y | - | T(7) | Arithmetic mean $\alpha = 1$ |
| Y | Y | Y | Y | Y | Y | - | - | Power mean $0 < \alpha < 1$ |
| Y | Y | Y | Y | Y | Y | Y | M(6) | Geometric mean $\alpha = 0$ |
| Y | Y | Y | Y | Y | Y | Y | - | Power mean $\alpha < 0$ |
| Y | Y | Y | Y | Y | - | Y | M(6) | Normalise by M(6), then $\alpha < 0$ |
| Y | Y | Y | Y | Y | - | - | T(7) | $\alpha = 2$ (Lin and Binns, 1988) |
| Y | Y | Y | Y | - | - | - | M(6), T(7), A(8) | Adaptability (St-Pierre et al., 1967) |
| Y | Y | Y | Y | - | - | - | All (6),(7),(8),(9) | TOP (Fox et al., 1990) |

*Notes:* We did not discuss the Adaptability rule of St-Pierre et al. (1967) in this paper. The rule ranks cultivars in accordance with their adaptability score, which is the percentage of environments where the cultivar's yield exceeds the environmental mean: $|\{j \in E : y_{ij} > \frac{1}{|C|} \sum_{k \in C} y_{kj}\}| * \frac{100}{|E|}$ .

Incomplete multi-environment trials require special attention. In practice, not all cultivars are evaluated in all environments, and some observations may be missing or removed because of experimental failure, disease damage, or data-quality concerns. Such incomplete structures are common in long-term and large-scale cultivar evaluation networks. As we mention above, normalisation is necessary in this scenario, however the impossibility result of Theorem 7 shows that no reasonable ranking rule can satisfy independence of irrelevant alternatives as well as a normalisation method. This should not be seen as a weakness of the proposed framework; rather, it exposes a fundamental limitation of ranking incomplete yield profiles without additional modelling assumptions.

For this reason, the proposed ranking rules should usually be applied to adjusted, estimated, or predicted yield profiles rather than directly to raw incomplete data. Statistical models can be used to estimate or predict cultivar yield across the relevant target environments. Once these yield profiles are obtained, the ranking rule can be applied as a separate decision step. This separation is important: statistical models estimate the entries of the yield profile, whereas the ranking rule determines how those entries are aggregated into a final recommendation.

From a practical breeding perspective, the framework suggests a transparent workflow. First, observe, estimate, or predict cultivar yield across environments using an appropriate statistical model. Second, define the target population of environments and specify environmental weights when some environments are more relevant than others. Third, choose and report the normalisation method, especially when environments differ substantially in productivity. Our analysis justifies multiplication normalisation. Fourth, rank cultivars using a power mean selected according to the desired level of risk aversion, with the geometric mean providing a natural default given that it produces a ranking that is consistent with multiplication normalisation. Finally, report the ranking rule, environmental weights, normalisation method, and treatment of missing, zero, or negative values so that cultivar recommendations are reproducible and interpretable.

In conclusion, the proposed axiomatic framework provides a principled basis for cultivar recommendation in multi-environment trials. It does not replace classical or modern methods for genotype-by-environment analysis. Instead, it complements them by clarifying the final decision step. Once cultivar yield has been observed, adjusted, estimated, or predicted, the breeder still needs a logically defensible rule for ranking cultivars. By making the assumptions behind that rule explicit, the framework helps ensure that cultivar recommendations are consistent with yield potential, protection against poor performance, environmental relevance, and the practical objectives of plant breeding.

## Appendix A. Additional Axioms and Theorems

Ranking theory is a very versatile formalism. In this paper, our focus has been on ranking cultivars based on yield data in a multi-environment trial, but the same mathematics can be used to describe sporting events (Kondratev et al., 2024; Jones and Wilson, 2025; Stival, 2025), average utility (Kothiyal et al., 2014), multi-criteria decisions (Wijnmalen and Wedley, 2009; Verly and De Smet, 2013), recommender systems (Christensen and Schiaffino, 2011), gene expression meta-analysis (Toro-Domínguez et al., 2021), or ensemble algorithms (Bolón-Canedo and Alonso-Betanzos, 2019). However, though the mathematics maybe universal, the choice of which properties are appropriate to a given setting depends very much on what we are trying to model.

In this section we will look at two properties – anonymity and strict separability – which are standard in the ranking literature (Wijnmalen and Wedley, 2009; Verly and De Smet, 2013; Kothiyal et al., 2014; Kondratev et al., 2024), but make little sense in the context of ranking cultivars.

The property of **anonymity**, informally, is satisfied by a ranking rule that treats all environments equally. Formally, a **permutation** of the identities of the environments is a one-to-one mapping *σ* between two sets of environments *E* and *E*^*′*^; we can treat 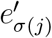 as a new identity of environment *e*_*j*_. For a permutation *σ* and a yield profile **Y** on *E*, we denote *σ*(**Y**) = **Y**^*′*^ the profile with permuted identities *E*^*′*^ of the environments, that is, 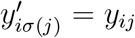 for all *i, j*. A ranking rule *R* satisfies anonymity if *R*(*σ*(**Y**)) = *R*(**Y**) for all *σ*, **Y**. In the case of the weighted power mean, the power mean satisfies anonymity if and only if all the weights are equal.

Anonymity is used to formalise notions of fairness or ignorance. In the case of an average utility, where the environments are the preferences of the voters, anonymity states that all voters are to be treated equally. In the case of an ensemble algorithm, where the environments are the solutions to a problem offered by various subroutines, anonymity states that we have no information about which subroutine is better at this stage, and we may as well give them equal weight.

**Strict separability** is about the effect of informationally “useless” environments on the final ranking. An environment *e*_*j*_ is said to be **indifferent** if *y*_*ij*_ = *y*_*kj*_ for all *i, k*: the yield of every cultivar in *e*_*j*_ is the same. A ranking rule satisfies strict separability if the removal of an indifferent environment leaves the ranking unchanged (Finan and Hurley, 2002).

Neither anonymity nor strict separability are appropriate for the context of ranking cultivars, and the main reason for this is that an environment in a multi-environment trial is not an atomic entity, but contains information about the year, the location, and the condition of the trial. Consider the yield profile in Table 5 where two cultivars were tested in six environments, over a period of four years (Y1, Y2, Y3, Y4) and in two locations (L1, L2). Note that the profile is not balanced: location L2 was not available in Y3, and L1 was not available in Y4.

**TABLE 5.** Unbalanced yield profile.

| Cultivar | Environment: yield (t/ha) |  |  |  |  |  |
| --- | --- | --- | --- | --- | --- | --- |
|  | Y1-L1 | Y1-L2 | Y2-L1 | Y2-L2 | Y3-L1 | Y4-L2 |
| $c_1$ | 9 | 8 | 8 | 9 | 9 | 9 |
| $c_2$ | 9 | 9 | 9 | 9 | 8 | 8 |

| C. | Yield grouped (t/ha) |  |  |  |  |  |
| --- | --- | --- | --- | --- | --- | --- |
|  | All data, L1, L2 | Two best years | Two worst years |  |  |  |
| $c_1$ | 9(67%) 8(33%) | 9(100%) | 9(50%) 8(50%) | | | |
| $c_2$ | 9(67%) 8(33%) | 9(100%) | **8(100%)** | | | |

If we know which years or locations are more relevant to how we plan to grow the cultivar, we can use this information to put more weight on those environments, violating anonymity. But if we do not, then we may reasonably assume that every year and every location is equally important. This sounds a bit like anonymity, and if we treat all environments equally we must conclude that the cultivars are equally good: after all, both have a yield of 9 four times and a yield of 8 twice. However, it may be more reasonable to treat every *year* and *location* equally, and if we look at the bottom half of Table 5 we see how each cultivar performs in its best/worst location and year. Note that the cultivars do the same everywhere, except in their two worst years, where *c*_1_ is better. So perhaps *c*_1_ is the better cultivar after all.

A similar argument will show us that in this context indifferent environments are not useless either. If we add two indifferent environments to the yield profile, Y3-L2 and Y4-L1, we get Table 6. Table 5 and Table 6 differ only by the addition of indifferent environments, so according to strict separability, the same cultivar should be chosen in both. However, even though we argued that *c*_1_ is better in Table 5 because it does better in more years, in Table 6 the two cultivars behave completely symmetrically, and should probably be ranked equally.

**TABLE 6.** Balanced yield profile with indifferent environments added.

| Cultivar | Environment: yield (t/ha) |  |  |  |  |  |  |  |
| --- | --- | --- | --- | --- | --- | --- | --- | --- |
|  | Y1-L1 | Y1-L2 | Y2-L1 | Y2-L2 | Y3-L1 | **Y3-L2** | **Y4-L1** | Y4-L2 |
| $c_1$ | 9 | 8 | 8 | 9 | 9 | **9** | **9** | 9 |
| $c_2$ | 9 | 9 | 9 | 9 | 8 | **9** | **9** | 8 |

*Notes:* The practical interpretation is the following: balanced yield profiles are recommended as input to ranking rules, even if some environments were informationally useless.

For these reasons, we keep the following theorems out of the main text. But for the curious, this is what we can say about anonymity and strict separability:

### Theorem 8.

Consider a domain with positive yields, complete yield profiles, and *fixed* sets of cultivars and environments with at least three cultivars. If a ranking rule satisfies multiplication invariance, Pareto efficiency, independence of irrelevant alternatives, separability, and anonymity, then it is a refinement of the minimum rule, maximum rule or power mean (Roberts, 1980, theorem 6).

In the case of *variable* sets of cultivars and environments, if a ranking rule satisfies multiplication invariance, Pareto efficiency, strict independence of irrelevant alternatives, separability, and anonymity, then for each *number* of environments, it is a refinement of the minimum rule, maximum rule, or power mean.

### Theorem 9.

1. if it also satisfies anonymity and admits multiplication normalisation (6), then it is a refinement of the geometric mean (follows from Moulin 1988, theorem 2.3).
2. if it also satisfies anonymity and admits translation normalisation (7), then it is a refinement of the arithmetic mean (follows from a corollary of theorem 8.1 in Bossert and Weymark 2004).

A ranking rule satisfies **no weak dictator** axiom if for each set of at least two cultivars and each set of at least two environments, no environment is a weak dictator. Anonymity implies no weak dictator axiom.

We say that the parameter *α*_*E*_ is **global** if it does not depend on the set of environments *E, α*_*E*_ = *α* for all *E*. We say that the weights of the weighted power mean are **global** if there exists a sequence of nonnegative numbers *w*_*j*_ such that

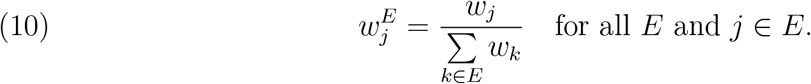

### Theorem 10.

Consider a domain with positive yields, complete yield profiles, and *variable* sets of cultivars and environments. If a ranking rule satisfies multiplication invariance, Pareto efficiency, strict independence of irrelevant alternatives, and strict separability, then:

1. if it also satisfies no weak dictator, then it is a refinement of the minimum rule, maximum rule, or the weighted power mean with global parameter *α* and positive global weights.
2. if it also satisfies reversibility, then it is a refinement of the minimum rule or the weighted power mean with global parameter *α* ≤ 0 and positive global weights.
3. if it also satisfies no weak dictator and admits translation normalisation (7), then it is a refinement of the weighted arithmetic mean with positive global weights.
4. if it also satisfies anonymity, then it is a refinement of the minimum rule, maximum rule, or power mean with global parameter *α*.

### Theorem 11.

Consider a domain with positive yields, complete yield profiles, and *variable* sets of cultivars and environments. If a ranking rule satisfies Pareto efficiency, strict independence of irrelevant alternatives, strict separability, one of reversibility or no weak dictator, and admits multiplication normalisation (6), then it is a refinement of the weighted geometric mean with positive global weights.

## Appendix B. Proofs of Theorems

### Theorem 2.

*Proof*. The theorem statement contains all the requirements of Theorem 1 plus reversibility. By Theorem 1, we know that any rule that is not a refinement of the maximum, minimum, or the weighted power mean does not satisfy this properties. Hence it is sufficient to construct counter examples for the maximum rule, and the weighted power mean with *α >* 1 or non-positive weights.

Consider a yield profile such that: *y*_*kj*_ = 1 for some *c*_*k*_ and for each *e*_*j*_; *y*_*ij*_ = *ε <* 1 for each *c*_*i*_≠ *c*_*k*_ and for each *e*_*j*_. The aggregate performance is 1 for *c*_*k*_ and *ε* for other cultivars.

If the rule is a refinement of the maximum rule and we downgrade *y*_*kj*_ for some *j*, then the aggregate performance of *c*_*k*_ is 1 and therefore *c*_*k*_ is ranked first, which contradicts reversibility. If the rule is a refinement of the weighted power mean where some *e*_*j*_ has a zero weight, and we downgrade *y*_*kj*_, then the aggregate performance of *c*_*k*_ is 1 and hence *c*_*k*_ is ranked first, which contradicts reversibility.

If the rule is a refinement of the weighted power mean, where all weights are positive, *w*_max_ ∈ (0, 1) is the maximum of these weights, and the parameter *α >* 0, then we choose *ε <* (1 − *w*_max_)^1*/α*^. If we downgrade *y*_*kj*_ for some *j*, then the aggregate performance of *c*_*k*_ is at least (1 − *w*_max_)^1*/α*^, and hence *c*_*k*_ is ranked first, which contradicts reversibility. □

Similarly, in Theorem 4, reversibility implies positive weights.

In Theorems 3, 4, 5, 8, 10, and 11 for variable size trials, if we strengthen IIA to SIIA, the same implications hold, as in the corresponding theorems for fixed size trials, but the parameters are replaced with 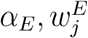. It is straightforward from the fact that the parameters for (*C, E*) and (*C*^*′*^, *E*) must be equivalent to the parameters for (*C* ∪ *C*^*′*^, *E*) and hence to each other.

### Theorem 6.

*Proof*. For (2), consider a yield profile **Y** with two cultivars and two environments where *y*_1, 1_ = 3, *y*_1, 2_ = 4, *y*_2, 1_ = 4, *y*_2, 2_ = 3. Without loss of generality, suppose *c*_1_ is not ranked last in the aggregate ranking; otherwise, the argument would be the same for *c*_2_. Reversibility requires that *c*_1_ be ranked last after some downgrade of the quantity *y*_1, 1_. However, any downgrade of *y*_1, 1_ does not change the ordinal ranking profile and the profile after affine normalisation.

For (1), for each *ε >* 0, denote **Y**_+*ε*_ the change of **Y** such that we only upgrade *y*_1, 1_ to 3 + *ε*. Due to monotonicity, *c*_1_ is not the last ranked in **Y**_+*ε*_. Denote 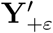 the change of **Y**_+*ε*_ where we only change *y*_1, 1_ to *ε* and *y*_2, 1_ to 1. Because **Y**_+*ε*_ and 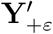 coincide after translation normalisation, *c*_1_ is not the last ranked in 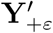 for each *ε >* 0, which contradicts reversibility. □

### Theorem 7.

*Proof*. Consider a yield profile with *c*_1_ and *c*_2_, whose yield was measured and used in disjoint sets of environments. By IIA, the relative ranking of *c*_1_ and *c*_2_ will remain intact if we change the measured and used yield of the other cultivars, making all environments almost indifferent. All such yield profiles coincide after normalisation and hence provide the same outcome. □

### Theorem 10.

*Proof*. The statements follow from the following fact. Suppose *C* = {*c*_1_, *c*_2_}, *E* = {*e*_*j*_, *e*_*k*_} and available ranking rules include only minimum rule, maximum rule, ranking by *e*_*j*_, ranking by *e*_*k*_, and weighted power means with positive weights. Then, for each pair of available rules, there exists a yield profile such that one rule ranks *c*_1_ above *c*_2_ and another rule ranks *c*_1_ below *c*_2_.

By Theorem 3, for each set of environments, the rule is a refinement of the minimum rule, maximum rule, or weighted power mean. For each set of two environments, any of the additional no weak dictator, anonymity, or reversibility excludes the possibility of zero weights. Let us define *w*_1_ = 1 and 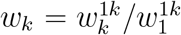 for all other *k*. By strict separability, for each pair of {*e*_*j*_, *e*_*k*_} and {*e*_*j*_, *e*_*k*_, *e*_*l*_}, both rules are refinements of the minimum rule, or both are refinements of the maximum rule, or both are refinements of the weighted power mean with *α*_*jk*_ = *α*_*jkl*_ and positive weights such that 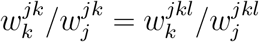.

For the latter case, substituting *l* = 1, we have

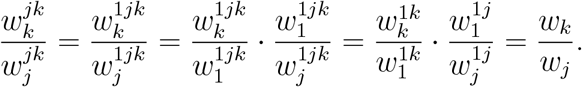

Hence, the weights 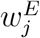 are proportional to *w*_*j*_ for all *E* with two or three environments. It is easy to see that *α*_12*j*_ = *α*_12_ = *α*_1*j*_ = *α*_2*j*_, *α*_1*jk*_ = *α*_1*j*_ = *α*_1*k*_ = *α*_*jk*_, *α*_2*jk*_ = *α*_2*j*_ = *α*_2*k*_ = *α*_*jk*_, *α*_*jkl*_ = *α*_*jk*_ = *α*_*jl*_ = *α*_*kl*_ and hence the parameters *α*_*E*_ are the same for all *E* with two or three environments. For each *E* of arbitrary size, we can consider any {*e*_*j*_, *e*_*k*_} with positive 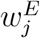. By strict separability, *α*_*jk*_ = *α*_*E*_ and 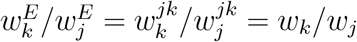

Items (1) and (2) follow from Theorem 3. Item (3) follows from Theorem 4. Item (4) follows from Theorem 8. □

The admission of multiplication normalisation implies multiplication invariance. Hence, Theorem 11 follows from Theorems 4 and 10.

## Appendix C. Pareto Inefficiency of Existing Methods

The main text motivates (weak) Pareto efficiency as a minimal consistency requirement for cultivar ranking: a cultivar that performs no worse than another cultivar in every environment should not be ranked below it. To stress how paradoxical a violation of Pareto efficiency is, recall that if a rule is not at least weakly Pareto efficient, then it is not monotonic – under such a ranking rule, increasing the yield of a cultivar could lead to that cultivar not being recommended.

In this appendix we extend that analysis by providing explicit counterexamples for many existing methods. The counterexamples cover several families of rules. We consider nonparametric rank-based stability measures, including Huhn’s *S*^(1)^, *S*^(2)^, *S*^(3)^, and *S*^(6)^ statistics and Thennarasu’s *NP* ^(1)^, *NP* ^(2)^, *NP* ^(3)^, and *NP* ^(4)^ statistics. We also consider variance-based stability measures, including Wricke’s ecovalence, Shukla’s stability variance, the mean variance component, the GE variance component, environmental variance, and the coefficient of variation. In addition, we study rank aggregation and selection rules, including Kang’s rank, the TOP rule, and average rank. Finally, we consider safety-first selection indices, AMMI-based indices, and the GGE biplot. Together, these examples show that Pareto inefficiency can arise across a wide range of standard methods for ranking cultivars in multi-environment trials.

We first introduce notation used throughout the appendix. Cultivar labels are specific to the profile under consideration. For the calculations below, let *m* = |*C*| and *n* = |*E*|.

We denote the mean yield of cultivar *i*, the mean yield in environment *j*, and the grand mean by

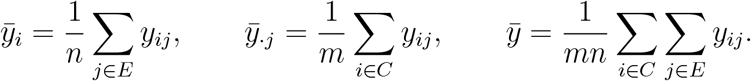

We denote environmental variance by

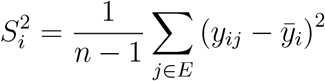

and environmental standard deviation by *S*_*i*_.

For rank-based measures, let *r*_*ij*_ be the rank of cultivar *i* in environment *j*, where rank 1 is best. We use inverse ranks

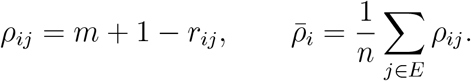

Larger values of *ρ*_*ij*_ are better. Let *M*_*di*_ be the median of *ρ*_*ij*_ across environments. For cultivar-centred yields, define

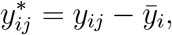

and let 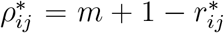 be the inverse rank of 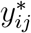 within environment *j*. We write 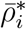 for the mean of 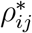 across environments and 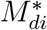 for its median.

In each counterexample, the Pareto-dominated cultivar is listed first, and the cultivar that dominates it is listed second. The values assigned by the corresponding rule to these two cultivars show the violation. The dominated cultivar receives a value at least as favorable as the value assigned to the cultivar that dominates it. Unless stated otherwise, smaller values are more favorable.

We begin with the profile from the main text, reported in Table 7.

**TABLE 7.** The profile from the main text.

| Cultivar | Environment: yield $y_{ij}$ (rank $r_{ij}$) | | | | | $\bar{y}_i$ (avg. rank) |
| --- | --- | --- | --- | --- | --- | --- |
| | $e_1$ | $e_2$ | $e_3$ | $e_4$ | $e_5$ | |
| $c_1$ | 13.32 (1) | 11.67 (1) | 11.07 (1) | 12.18 (2) | 11.08 (2) | 11.86 (1.4) |
| $c_2$ | 12.39 (2) | 10.89 (4) | 10.65 (4) | 12.56 (1) | 11.65 (1) | 11.63 (2.4) |
| $c_3$ | 11.48 (3) | 11.59 (2) | 10.73 (3) | 12.14 (3) | 11.01 (3) | 11.39 (2.8) |
| $c_4$ | 10.92 (5) | 11.52 (3) | 10.92 (2) | 11.25 (5) | 10.49 (5) | 11.02 (4.0) |
| $c_5$ | 11.26 (4) | 10.56 (5) | 10.60 (5) | 11.46 (4) | 10.65 (4) | 10.91 (4.4) |
| $c_6$ | 10.68 (6) | 10.48 (6) | 10.32 (6) | 11.24 (6) | 10.19 (6) | 10.58 (6.0) |
| $c_7$ | 10.43 (7) | 9.75 (7) | 8.25 (9) | 10.43 (7) | 9.75 (7) | 9.72 (7.4) |
| $c_8$ | 9.95 (8) | 8.46 (8) | 8.86 (7) | 10.21 (8) | 9.37 (8) | 9.37 (7.8) |
| $c_9$ | 9.86 (9) | 8.28 (9) | 7.16 (10) | 9.63 (9) | 8.18 (9) | 8.62 (9.2) |
| $c_{10}$ | 9.58 (10) | 7.18 (10) | 8.27 (8) | 9.36 (10) | 7.41 (10) | 8.36 (9.6) |
| $\bar{y}_{\cdot j}$ | 10.99 | 10.04 | 9.68 | 11.05 | 9.98 | $\bar{y}=10.35$ |

Table 8 reports the cultivar-centred yields 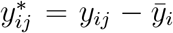. The values in parentheses are the corresponding inverse ranks 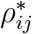.

**TABLE 8.**
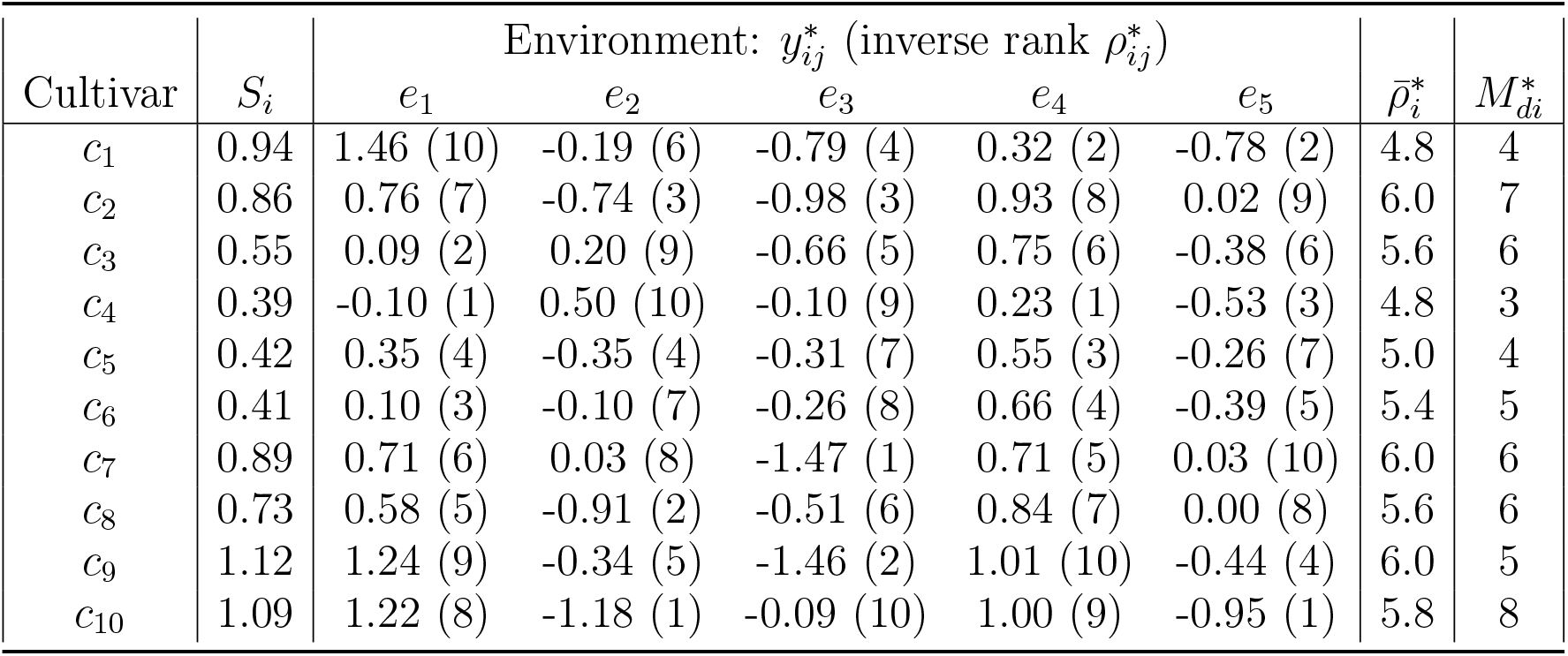
Preliminary calculations for the profile from the main text.

**TABLE 9.** Wricke Ecovalence.

| Cultivar | $c_1$ | $c_2$ | $c_3$ | $c_4$ | $c_5$ | $c_6$ | $c_7$ | $c_8$ | $c_9$ | $c_{10}$ | $\sum W_i^2$ |
| --- | --- | --- | --- | --- | --- | --- | --- | --- | --- | --- | --- |
| $W_i^2$ | 1.01 | 0.5 | 0.56 | 1.77 | 0.25 | 0.5 | 0.93 | 0.54 | 1.1 | 1.85 | 9.02 |

**TABLE 10.** Stability variance.

| Cultivar | $\sigma_i^2$ (stability variance rank) |
| --- | --- |
| $c_1$ | 0.2858 (7) |
| $c_2$ | 0.1264 (3) |
| $c_3$ | 0.1450 (5) |
| $c_4$ | 0.5206 (9) |
| $c_5$ | 0.0453 (1) |
| $c_6$ | 0.1251 (2) |
| $c_7$ | 0.2589 (6) |
| $c_8$ | 0.1389 (4) |
| $c_9$ | 0.3113 (8) |
| $c_{10}$ | 0.5476 (10) |

The following subsections give the definitions and calculations for the counterexamples based on Tables 7 and 8.

### C.1. *S*^(1)^ statistic

It measures the average absolute difference in inverse ranks across pairs of environments (Hühn, 1979):

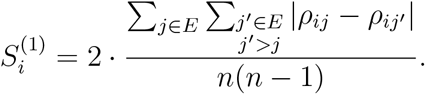

- For 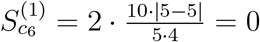
- For 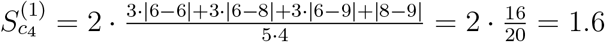

### C.2. *S*^(2)^ statistic

It measures the variance of inverse ranks around the mean inverse rank (Hühn, 1979):

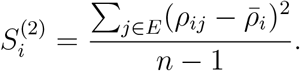

- For 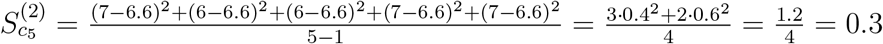
- For 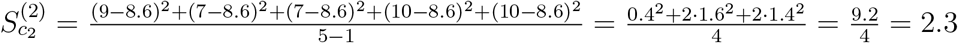

### C.3. *S*^(3)^ statistic

It measures squared inverse-rank deviations, normalized by the mean inverse rank (Hühn, 1979):

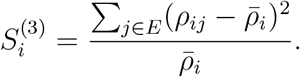

- For 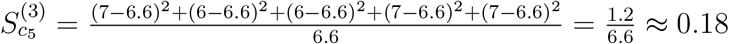
- For 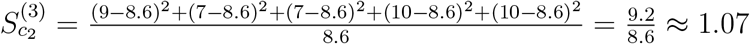

### C.4. *S*^(6)^ statistic

It measures absolute inverse-rank deviations, normalized by the mean inverse rank (Hühn, 1979):

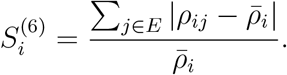

- For 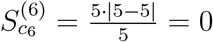
- For 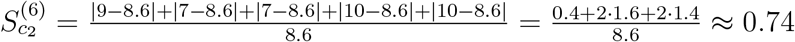

### C.5. *NP* ^(1)^ statistic

It measures the average absolute difference between adjusted ranks and the adjusted median rank (Thennarasu, 1995):

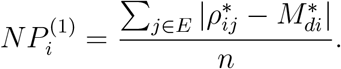

- For 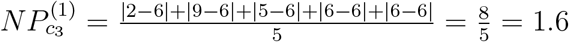
- For 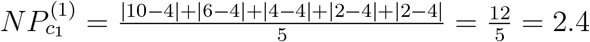

### C.6. *NP* ^(2)^ statistic

It measures the deviation of adjusted ranks from the adjusted median rank relative to the unadjusted median rank (Thennarasu, 1995):

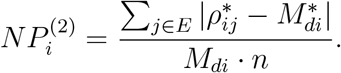

- For 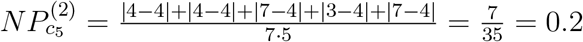
- For 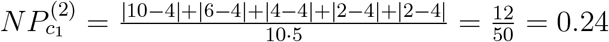

### C.7. *NP* ^(3)^ statistic

It is defined as the normalized standard deviation of adjusted ranks (Thennarasu, 1995):

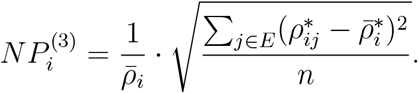

- For 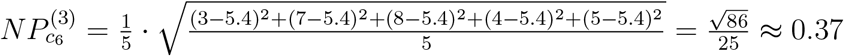
- For 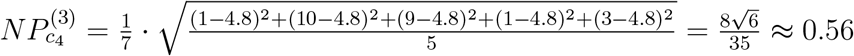

### C.8. *NP* ^(4)^ statistic

It measures the average absolute difference between pairs of adjusted ranks, scaled by the mean unadjusted rank (Thennarasu, 1995):

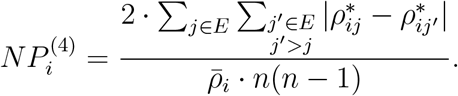

- For 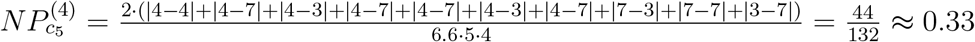
- For 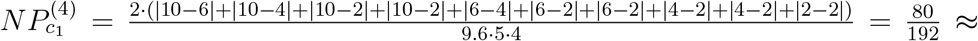

### C.9. Wricke ecovalence

It measures the contribution of the cultivar to genotype-by-environment interaction (Wricke, 1962):

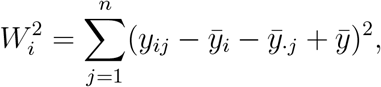

where 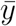 is the grand mean of the yield.

- For 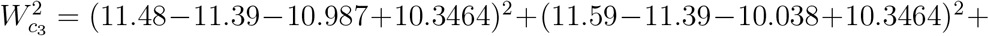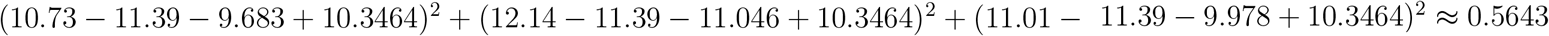
- For 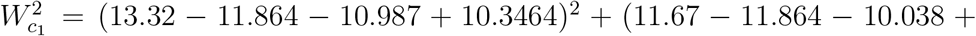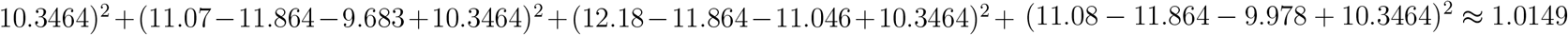

### C.10. Shukla’s stability variance

It adjusts Wricke ecovalence by a common term based on total ecovalence (Shukla, 1972):

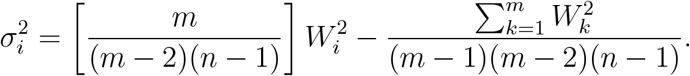

Wricke Ecovalence has been calculated analogously for the remaining cultivars. The rounded results are presented in the following table:

- For 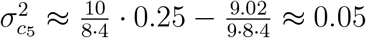
- For 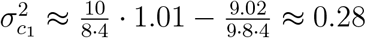

### C.11. Kang’s rank

It uses both mean yield and Shukla’s stability variance to identify high-yielding and stable cultivars (Kang, 1988):

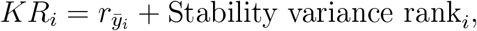

where 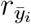 is the rank based on 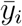 and the Stability variance rank is based on 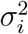.

- For 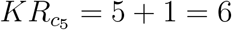
- For 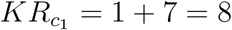

Table 11 reports a profile used for an additional Kang’s-rank counterexample.

**TABLE 11.** Kang’s-rank counterexample profile.

| Cultivar | Env.: yield $y_{ij}$ | | $\bar{y}_i$ |
| --- | --- | --- | --- |
| | $e_1$ | $e_2$ | |
| $b_1$ | 13.1 | 11.1 | 12.1 |
| $b_2$ | 10.0 | 11.0 | 10.5 |
| $b_3$ | 9.0 | 10.0 | 9.5 |
| $\bar{y}_{\cdot j}$ | 10.7 | 10.7 | 10.7 |

**TABLE 12.** Stability variance: Kang’s-rank counterexample.

| Cultivar | $\sigma_i^2$ (stability variance rank) |
| --- | --- |
| $b_1$ | 4.5 (3) |
| $b_2$ | 0 (1.5) |
| $b_3$ | 0 (1.5) |

For Table 11, the Wricke ecovalences are

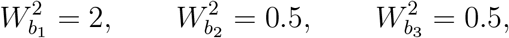

and

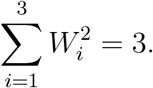

Since *m* = 3 and *n* = 2,

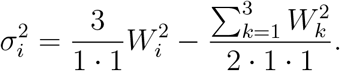

This gives the following table:

- For 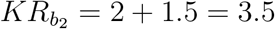
- For 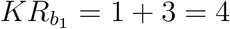

### C.12. Mean variance component

It measures a cultivar’s stability through its average contribution to genotype-by-environment interaction variance (Plaisted and Peterson, 1959):

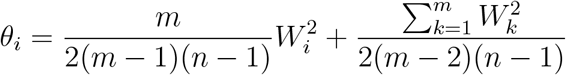

- For 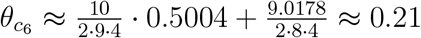
- For 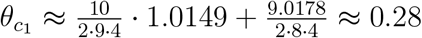

### C.13. GE variance component

It is a modified measure in which higher values indicate greater stability (Plaisted, 1960):

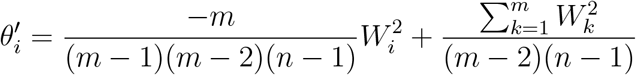

For this rule, larger values are better.

- For 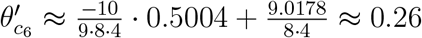
- For 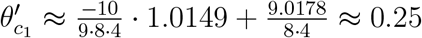

### C.14. Environmental variance

It measures stability through the variance of yield across environments (Römer, 1917):

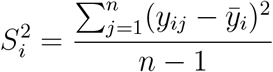

- For 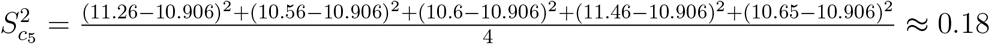
- For 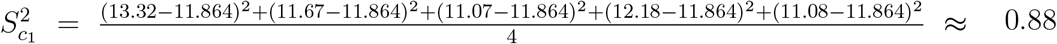

### C.15. Coefficient of variation

It measures the relative variability of yield across environments as a percentage of mean yield (Francis and Kannenberg, 1978):

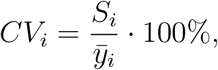

- For 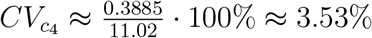
- For 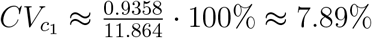

### C.16. TOP rule

The TOP rule ranks cultivars by the share of environments in which a cultivar belongs to the top third of cultivars (Fox et al., 1990):

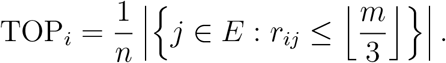

For TOP, larger values are better.

In Table 7, the top third consists of cultivars with ranks at most ⌊10*/*3⌋ = 3:

- For 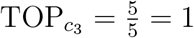
- For 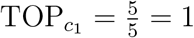

### C.17. Average rank

It jointly evaluates cultivar performance and stability using the average rank (AR) across yield and multiple statistics, following the approach of Pour-Aboughadareh et al. (2019):

The tables below report rounded values. The ranks in parentheses are computed from unrounded values.

**TABLE 13.**
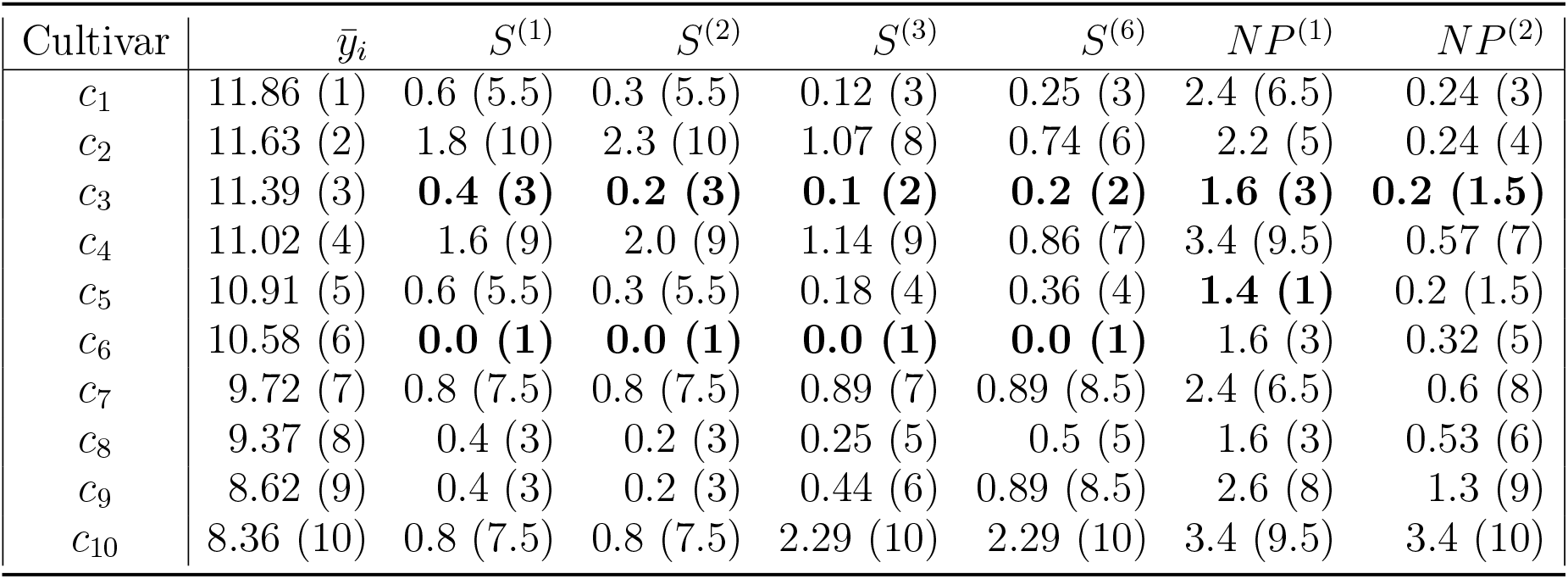
Performance metrics for the main example: part 1 (out of 3)

| Cultivar | $\bar{y}_i$ | $S^{(1)}$ | $S^{(2)}$ | $S^{(3)}$ | $S^{(6)}$ | $NP^{(1)}$ | $NP^{(2)}$ |
| --- | --- | --- | --- | --- | --- | --- | --- |
| $c_1$ | 11.86 (1) | 0.6 (5.5) | 0.3 (5.5) | 0.12 (3) | 0.25 (3) | 2.4 (6.5) | 0.24 (3) |
| $c_2$ | 11.63 (2) | 1.8 (10) | 2.3 (10) | 1.07 (8) | 0.74 (6) | 2.2 (5) | 0.24 (4) |
| $c_3$ | 11.39 (3) | **0.4 (3)** | **0.2 (3)** | **0.1 (2)** | **0.2 (2)** | **1.6 (3)** | **0.2 (1.5)** |
| $c_4$ | 11.02 (4) | 1.6 (9) | 2.0 (9) | 1.14 (9) | 0.86 (7) | 3.4 (9.5) | 0.57 (7) |
| $c_5$ | 10.91 (5) | 0.6 (5.5) | 0.3 (5.5) | 0.18 (4) | 0.36 (4) | **1.4 (1)** | 0.2 (1.5) |
| $c_6$ | 10.58 (6) | **0.0 (1)** | **0.0 (1)** | **0.0 (1)** | **0.0 (1)** | 1.6 (3) | 0.32 (5) |
| $c_7$ | 9.72 (7) | 0.8 (7.5) | 0.8 (7.5) | 0.89 (7) | 0.89 (8.5) | 2.4 (6.5) | 0.6 (8) |
| $c_8$ | 9.37 (8) | 0.4 (3) | 0.2 (3) | 0.25 (5) | 0.5 (5) | 1.6 (3) | 0.53 (6) |
| $c_9$ | 8.62 (9) | 0.4 (3) | 0.2 (3) | 0.44 (6) | 0.89 (8.5) | 2.6 (8) | 1.3 (9) |
| $c_{10}$ | 8.36 (10) | 0.8 (7.5) | 0.8 (7.5) | 2.29 (10) | 2.29 (10) | 3.4 (9.5) | 3.4 (10) |

- For 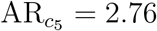
- For 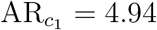

**TABLE 14.** Performance metrics for the main example: part 2 (out of 3)

| Cultivar | $NP^{(3)}$ | $NP^{(4)}$ | $W_i^2$ | $\sigma_i^2$ | KR | $\theta_i$ | $\theta'_i$ |
| --- | --- | --- | --- | --- | --- | --- | --- |
| $c_1$ | 0.31 (4) | 0.42 (4) | 1.01 (7) | 0.29 (7) | 8.0 (4) | 0.28 (7) | 0.25 (7) |
| $c_2$ | 0.29 (3) | 0.4 (3) | 0.5 (3) | 0.13 (3) | 5.0 (1) | 0.21 (3) | 0.26 (3) |
| $c_3$ | **0.27 (2)** | **0.37 (2)** | 0.56 (5) | 0.15 (5) | 8.0 (4) | 0.22 (5) | 0.26 (5) |
| $c_4$ | 0.56 (6) | 0.74 (6) | 1.77 (9) | 0.52 (9) | 13.0 (7.5) | 0.39 (9) | 0.22 (9) |
| $c_5$ | **0.25 (1)** | **0.33 (1)** | **0.25 (1)** | **0.05 (1)** | 6.0 (2) | **0.17 (1)** | **0.27 (1)** |
| $c_6$ | 0.37 (5) | 0.52 (5) | 0.5 (2) | 0.13 (2) | 8.0 (4) | 0.21 (2) | 0.26 (2) |
| $c_7$ | 0.84 (8) | 1.17 (8) | 0.93 (6) | 0.26 (6) | 13.0 (7.5) | 0.27 (6) | 0.25 (6) |
| $c_8$ | 0.64 (7) | 0.88 (7) | 0.54 (4) | 0.14 (4) | 12.0 (6) | 0.22 (4) | 0.26 (4) |
| $c_9$ | 1.69 (9) | 2.33 (9) | 1.1 (8) | 0.31 (8) | 17.0 (9) | 0.29 (8) | 0.24 (8) |
| $c_{10}$ | 2.84 (10) | 3.71 (10) | 1.85 (10) | 0.55 (10) | 20.0 (10) | 0.4 (10) | 0.22 (10) |

**TABLE 15.** Performance metrics for the main example: part 3 (final)

| Cultivar | $S_i^2$ | $CV_i$ | TOP | AR |
| --- | --- | --- | --- | --- |
| $c_1$ | 0.88 (8) | 7.89 (7) | 1.0 (1.5) | 4.94 (5) |
| $c_2$ | 0.74 (6) | 7.39 (5) | 0.6 (3) | 4.59 (4) |
| $c_3$ | **0.3 (4)** | **4.79 (4)** | 1.0 (1.5) | **3.24 (3)** |
| $c_4$ | **0.15 (1)** | **3.53 (1)** | 0.4 (4) | 6.82 (7) |
| $c_5$ | 0.18 (3) | 3.87 (2) | 0.0 (7.5) | **2.76 (1)** |
| $c_6$ | 0.17 (2) | 3.88 (3) | 0.0 (7.5) | 3.09 (2) |
| $c_7$ | 0.79 (7) | 9.16 (8) | 0.0 (7.5) | 7.18 (8) |
| $c_8$ | 0.53 (5) | 7.79 (6) | 0.0 (7.5) | 5.15 (6) |
| $c_9$ | 1.25 (10) | 12.97 (9) | 0.0 (7.5) | 7.76 (9) |
| $c_{10}$ | 1.2 (9) | 13.09 (10) | 0.0 (7.5) | 9.47 (10) |

The bold entries in the preceding tables highlight the main implication. The frontier obtained from mean yield and a stability statistic can include cultivars that are Pareto-dominated in the original yield profile. Hence, summarizing performance by mean yield and stability can fail to preserve Pareto efficiency.

### C.18. Safety-First Selection Indices

Table 16 reports the additional profile used for three of the safety-first counterexamples.

**TABLE 16.** Safety-first counterexample profile.

| Cultivar | Environ.: yield $y_{ij}$ | | | $\bar{y}_i$ |
| --- | --- | --- | --- | --- |
| | $e_1$ | $e_2$ | $e_3$ | |
| $a_1$ | 11.01 | 12.50 | 9.75 | 11.09 |
| $a_2$ | 10.56 | 11.35 | 11.32 | 11.08 |
| $a_3$ | 10.52 | 10.96 | 9.71 | 10.40 |
| $a_4$ | 10.20 | 9.45 | 8.31 | 9.32 |
| $\bar{y}_{\cdot j}$ | 10.57 | 11.07 | 9.77 | 10.47 |

These indices combine mean yield with a stability penalty (Eskridge, 1990). The general form is:

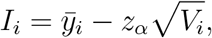

where *z*_*α*_ is a standard normal quantile and *V*_*i*_ is a stability measure for cultivar *i*.

The counterexample for 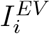 uses Table 7. The counterexamples for 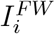, 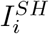, and 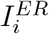 use Table 16. For the safety-first profile, the Shapiro-Wilk test does not reject normality for any cultivar at the 5% level. We use the one-sided normal quantile *z*_*α*_ = 1.645 at the 5% level throughout. For all safety-first indices, larger values are better.

#### C.18.1. Variance across environments

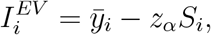

- For *c*_3_:

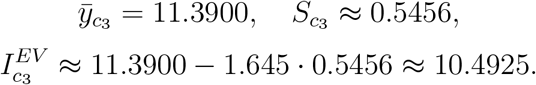
- For *c*_1_:

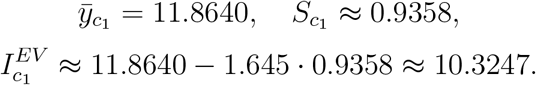

#### C.18.2. Finlay-Wilkinson regression

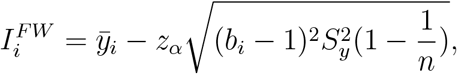

where 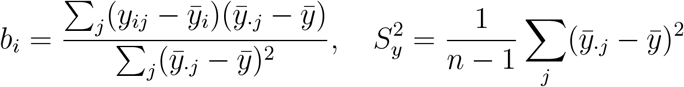

The cultivar means and regression coefficients (rounded to 4 decimals) are:

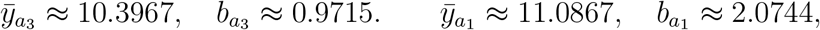

with *n* = 3 and 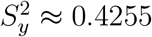

- For *a*_3_:

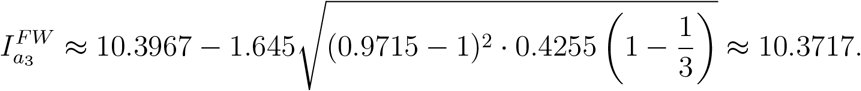
- For *a*_1_:

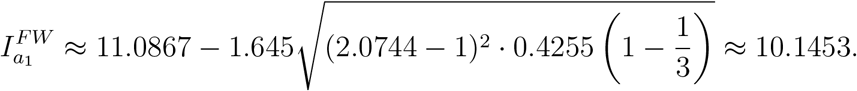

#### C.18.3. Shukla’s stability variance

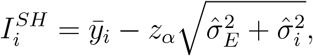

Where 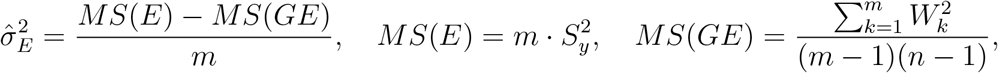,

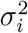 is Shukla’s stability variance, and 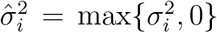 is the truncated value used in the index.

Wricke ecovalence 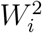 and Shukla’s stability variance 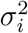 have been calculated for all cultivars. The results (rounded to 4 decimals) are:

**TABLE 17.** Wricke ecovalence and Shukla’s stability variance.

| Cultivar | $W_i^2$ | $\sigma_i^2$ |
| --- | --- | --- |
| $a_1$ | 1.1103 | 0.8268 |
| $a_2$ | 1.3720 | 1.0885 |
| $a_3$ | 0.0016 | -0.2820 |
| $a_4$ | 0.9184 | 0.6349 |

Negative estimates of 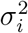 (such as for *a*_3_) are usually truncated to zero in the safety-first index.

The environmental component is

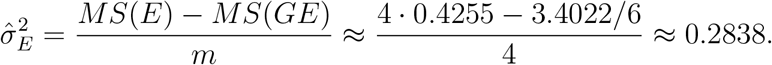

1. For *a*_3_:

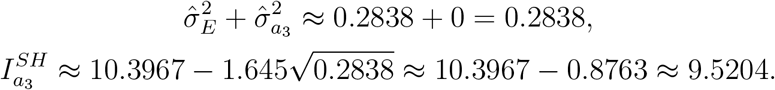
2. For *a*_1_:

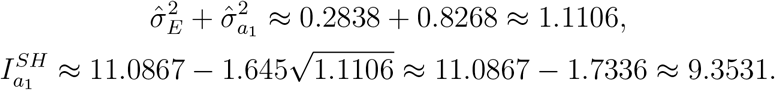

#### C.18.4. Eberhart-Russell model

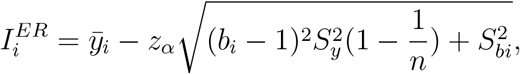

where 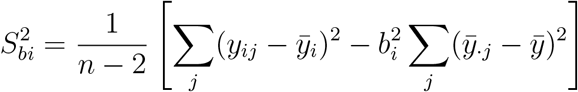

For the cultivars of interest, the relevant quantities (rounded to 4 decimals) are

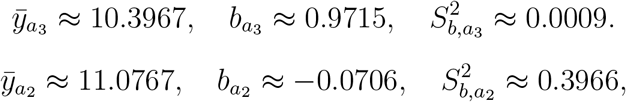

with *n* = 3 and 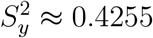

For *a*_3_:

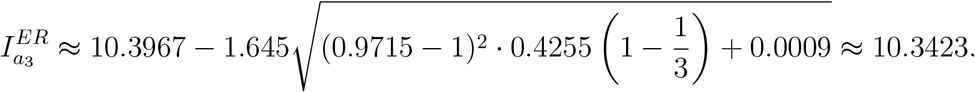

For *a*_2_:

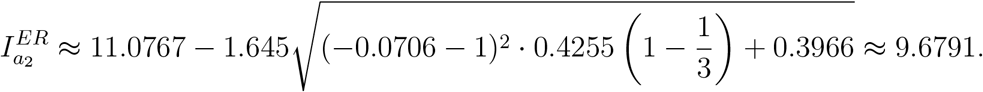

### C.19. AMMI-based indices

The following replicated profile is used for the AMMI counterexamples.

For the AMMI profile, we use two replications in each environment. A replication is a repeated observation of the same cultivar-environment pair. We denote by *y*_*ijk*_ the yield of cultivar *i* in environment *j* in replication *k*. The column *e*_*j, k*_ corresponds to environment *j* and replication *k*.

**TABLE 18.** AMMI counterexample profile.

| Cultivar | Environment-replication cell: yield $y_{ijk}$ | | | | | | | |
| --- | --- | --- | --- | --- | --- | --- | --- | --- |
| | $e_{1,1}$ | $e_{1,2}$ | $e_{2,1}$ | $e_{2,2}$ | $e_{3,1}$ | $e_{3,2}$ | $e_{4,1}$ | $e_{4,2}$ |
| $g_1$ | 11.11 | 11.44 | 10.67 | 10.18 | 13.82 | 13.41 | 11.06 | 11.74 |
| $g_2$ | 9.31 | 10.25 | 10.38 | 10.99 | 10.45 | 11.82 | 12.36 | 13.1 |
| $g_3$ | 9.85 | 10.66 | 9.39 | 10.47 | 10.2 | 10.68 | 11.72 | 12.84 |
| $g_4$ | 10.43 | 9.51 | 9.43 | 10.81 | 10.18 | 11.04 | 11.69 | 11.83 |
| $g_5$ | 8.4 | 9.23 | 11.02 | 11.81 | 10.02 | 10.7 | 10.26 | 11.03 |
| $g_6$ | 9.71 | 9.73 | 9.12 | 8.95 | 12.04 | 11.31 | 10.05 | 10.49 |
| $g_7$ | 6.71 | 8.38 | 8.41 | 7.76 | 9.12 | 9.6 | 9.5 | 10.02 |

The profile satisfies the standard preliminary checks for AMMI analysis. The genotype-by-environment interaction is significant at the 5% level according to the joint ANOVA *F*-test. The profile has no missing values or outliers, and the Shapiro-Wilk and Bartlett tests do not reject residual normality and homogeneity of variances at the 5% level.

C.19.1. *AMMI model*. The Additive Main Effects and Multiplicative Interaction (AMMI) model (Zobel et al., 1988) combines additive main effects with a multiplicative decomposition of genotype-by-environment interaction. For cultivar *i* in environment *j* and replication *k*, the model is:

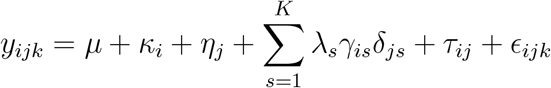

where *µ* is the grand mean, *κ*_*i*_ is the main effect of cultivar *i, η*_*j*_ is the main effect of environment *j, λ*_*s*_ is the eigenvalue of the *s*th interaction PCA (IPCA) retained in the model, and *γ*_*is*_ and *δ*_*js*_ are the corresponding cultivar and environment eigenvectors. Here *K* is the number of retained IPCAs, *τ*_*ij*_ is the GEI residual, and *ϵ*_*ijk*_ is the random error.

AMMI analysis was performed using the agricolae R package (de Mendiburu and Yaseen, 2020). Its outputs were processed with the ammistability R package (Ajay et al., 2023) to compute the stability statistics described in Pour-Aboughadareh et al. (2022). The notation in the following tables follows the output of the ammistability package. Since we are interested in both stability and yield, we consider Simultaneous Selection Indices (SSI) rather than the stability statistics alone.

The Yield Stability Index (YSI) (Farshadfar et al., 2011) is

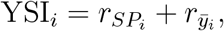

where 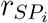 is the rank of the cultivar’s stability measure and 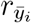 is the rank of the cultivar’s mean yield.

The next two tables show that all YSI-based indices rank *g*_7_ above *g*_5_, although *g*_5_ Paretodominates *g*_7_.

**TABLE 19.** Yield Stability Index-based indices: part 1 (out of 2)

| Cultivar | AMGE | ASI | ASV | ASTAB | AVAMGE | DA | DZ | EV | FA | MASI |
| --- | --- | --- | --- | --- | --- | --- | --- | --- | --- | --- |
| $g_7$ | 10 | 8 | 8 | 8 | 8 | 8 | 8 | 8 | 8 | 8 |
| $g_5$ | 12 | 10 | 10 | 12 | 11 | 11 | 12 | 12 | 11 | 10 |

**TABLE 20.** Yield Stability Index-based indices: part 2 (final)

| Cultivar | MASV | SIPC | Za |
| --- | --- | --- | --- |
| $g_7$ | 8 | 8 | 8 |
| $g_5$ | 10 | 12 | 11 |

The SSI of Rao and Prabhakaran (2005) is

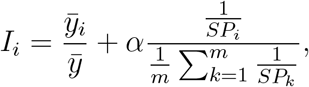

where *SP*_*i*_ is the stability measure of cultivar *i* and *α* = *w*_2_*/w*_1_, with *w*_1_ + *w*_2_ = 1. Higher values of *I*_*i*_ are better. In this example, *α* = 0.05.

The next two tables show that all SSI-based indices rank *g*_7_ above *g*_6_, although *g*_6_ Paretodominates *g*_7_.

**TABLE 21.** SSI-based indices: part 1 (out of 2)

| Cultivar | AMGE | ASI | ASV | ASTAB | AVAMGE | DA | DZ | EV | FA | MASI |
| --- | --- | --- | --- | --- | --- | --- | --- | --- | --- | --- |
| $g_7$ | 1.08 | 1.04 | 1.04 | 1.14 | 1.03 | 1.03 | 1.02 | 1.13 | 1.14 | 1.04 |
| $g_6$ | 0.96 | 0.99 | 0.99 | 0.98 | 0.99 | 0.99 | 1.00 | 0.98 | 0.98 | 0.99 |

**TABLE 22.** SSI-based indices: part 2 (final)

| Cultivar | MASV | SIPC | Za |
| --- | --- | --- | --- |
| $g_7$ | 1.04 | 1.02 | 1.03 |
| $g_6$ | 0.99 | 1.00 | 1.00 |

### C.20. GGE Biplot

Table 23 reports the profile used for the GGE biplot counterexample.

**TABLE 23.** GGE biplot counterexample profile.

| Cultivar | Env.: yield $y_{ij}$ | |
| --- | --- | --- |
| | $e_1$ | $e_2$ |
| $g_1$ | 10.57 | 7.5 |
| $g_2$ | 8.85 | 9.13 |
| $g_3$ | 9.92 | 7.34 |

The GGE biplot is a graphical tool for visualising cultivar effects and genotype-by-environment interaction in multi-environment trials (Yan and Tinker, 2006). In the cultivar-ranking view, cultivars are evaluated by their projection on the Average Environment Axis (AEA), which represents mean performance, and their deviation from the AEA, which represents instability. The ideal cultivar has the largest projection on the AEA and no deviation from it.

For the profile in Table 23, *g*_3_ is Pareto-dominated by *g*_1_, but is closer to the ideal point than *g*_1_.

**Fig. 1.**
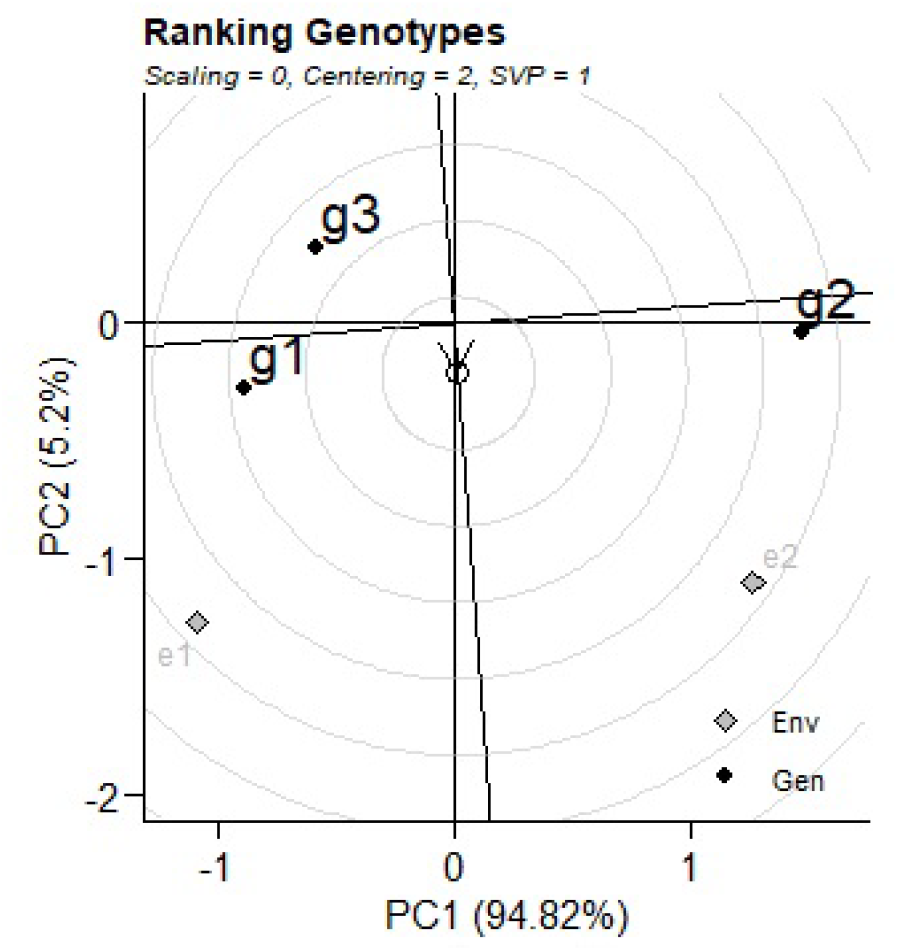
The biplot was computed using the metan R package (Olivoto and Lúcio, 2020) with environment-centred data, no scaling, and genotype-metric preserving singular value partitioning

## Footnotes

1 Reversibility is almost identical to strong reversal in Fortemps and Pirlot (2004) and similar to annihilator in Grabisch et al. (2011).

